# Structural brain alterations in autism: A voxel-based morphometry mega-analysis in 3,051 participants across 51 sites

**DOI:** 10.64898/2026.07.03.736397

**Authors:** Priya Rajagopalan, Gaon S. Kim, L. Nate Overholtzer, Emily Laltoo, Emma Gleave, Sebastian M. Benavidez, Chloe Retika, Paul M. Thompson, Katherine E. Lawrence

**Affiliations:** Imaging Genetics Center, Mark and Mary Stevens Neuroimaging and Informatics Institute, University of Southern California, Los Angeles, CA, USA; Department of Radiology, Los Angeles General Medical Center, Los Angeles, CA, USA; USC-Caltech MD-PhD Program, Keck School of Medicine of USC, Los Angeles, CA, USA; Department of Psychiatry and Biobehavioral Science, University of California Los Angeles, Los Angeles, CA, USA

**Keywords:** autism, voxel-based morphometry, gray matter volume, white matter volume, brain structure

## Abstract

**Background:** Previous large-scale structural MRI analyses of the brain in autism have identified gray matter (GM) differences when using region-of-interest analyses based on gross anatomical regions. However, such analyses have limited spatial specificity and may obscure subtle focal differences. Whole brain voxel-based morphometry (VBM) analyses enable greater spatial precision to identify and localize previously undetected neuroanatomical alterations.

**Purpose:** To rigorously identify voxel-wise GM and white matter (WM) volume differences in autism in the largest VBM mega-analysis to date.

**Materials and Methods:** This retrospective mega-analysis included structural 3D volumetric T1-weighted MRI brain scans from 3,051 participants (15.0 ± 8.2 yrs; 76.8% male; 1,519 autism; 1,532 neurotypicals) collected across 51 sites/scanners. Voxel-wise GM and WM volumes were quantified using the ENIGMA CAT12 VBM pipeline. Linear mixed-effects regression was performed at each voxel to evaluate the association between diagnostic group and voxel-wise volume while adjusting for standard nuisance covariates

**Results:** Autism was associated with widespread lower GM volume involving cortical, subcortical, and cerebellar regions (peak t=7.39, peak *β*=0.13); such GM differences were most notably detected in the bilateral orbitofrontal cortex, amygdala, thalamus, and posterior lobes of the cerebellum. WM volume was lower in autism across major projection, commissural, association, and cerebellar/brainstem tracts (peak t=6.74, peak *β*=0.08), including the *corona radiata*, internal capsule, corpus callosum, and cerebellar peduncles. These findings remained consistent in sensitivity analyses, including covarying for full-scale IQ and the application of increasingly strict motion exclusion criteria.

**Conclusion:** Autism is associated with smaller voxel-wise GM and WM volume involving widespread cortical, subcortical, and cerebellar regions. This high-resolution identification and localization of structural brain differences support the involvement of distributed neural systems in autism that underlie reward processing, sensory integration, and motor functioning in autism.

**Summary statement:** In the largest voxel-based morphometry study of autism to date, widespread smaller gray and white matter volumes were identified across distributed brain regions implicated in reward and sensorimotor function.

**Key Results:**

- Our largest voxel-based morphometry mega-analysis to date in autism (N=3,051), identified widespread gray and white matter alterations (peak t=7.39, peak *β*=0.13).
- We reveal novel associations within thalamic mediodorsal nuclei by leveraging finer voxel-wise volumetric analyses.
- Additional associations in orbitofrontal cortex, amygdala, and cerebellum, along with extensive white matter alterations across projection, commissural, association fibers, reinforce reward-processing, sensory and motor development theories in autism.

## Introduction

Autism is a neurodevelopmental condition that affects approximately 1 in 127 people worldwide.^1^ It is characterized by differences in social communication, alongside restricted and repetitive behaviors. Understanding the neuroanatomical differences associated with autism will be important for elucidating the neurobiological basis of its clinical manifestations.

The largest harmonized structural MRI analyses of autism so far used atlas-based gross anatomical region-of-interest (ROI) parcellations and reported lower subcortical volumes, increased frontal cortical thickness, and decreased temporal cortical thickness in autism.^2^ The study established an important foundation for understanding autism-related brain structure but inherently summarized biologically distinct subregions within relatively large anatomical parcels.^3^ To identify nuanced brain alterations in autism that may be obscured by averaging within anatomically defined regions, approaches with finer spatial resolution are needed for more precise localization.

Whole-brain voxel-based morphometry (VBM) is a method that provides greater spatial sensitivity than atlas-based ROI analyses, by allowing for the identification and localization of even spatially focal volumetric differences. However, VBM literature in autism remains limited. The largest multi-site VBM study (N=833) found that, on average, autism was associated with greater gray matter (GM) volume in the superior temporal gyrus.^4^ Yet, subsequent VBM meta-analyses have revealed inconsistent results. One study noted smaller GM volumes in autism in the amygdala, parahippocampal gyrus, and putamen.^5^ Another study reported smaller GM volumes in autism in the anterior cingulate cortex, medial prefrontal cortex, and cerebellum, as well as greater GM volumes in the middle temporal gyrus, olfactory cortex, and precentral gyrus.^6^ These inconsistencies across studies may reflect inadequate statistical power from extensive voxel-wise multiple comparisons, methodological heterogeneity including variability in processing pipelines, and/or limitations inherent to coordinate-based meta-analysis, which typically only consider published peak coordinates.^7^ Previous VBM studies focused almost entirely on GM, despite evidence from other modalities of widespread white matter (WM) involvement.^6,8^ Consequently, there remains a need for large-scale voxel-wise mega-analyses using harmonized processing pipelines that examine both GM and WM across the whole brain.

Here, we performed the largest VBM mega-analysis of GM and WM in autism to date (N=3,051), integrating structural brain MRI from 51 sites/scanners using a harmonized pipeline. By combining the statistical power of a large-scale sample with the spatial specificity of whole-brain voxel-wise analyses, we hypothesized that we would extend previous ROI-based findings by identifying novel, focal differences in previously undetected gray and white matter regions.

## Methods

### Participants and MRI Acquisition

This study is a retrospective mega-analysis of prospectively acquired data collected from 51 sites/scanners. Eligible participant data were obtained from datasets from the NIMH Data Archive (NDA), the Autism Brain Imaging Data Exchange (ABIDE), and the Healthy Brain Network (HBN) to increase generalizability by including data from many different samples.^9,10^ Demographic and scanner information from the contributing sites is summarized in **Table S1-2**. Briefly, structural T1-weighted (T1w) 3D volumetric brain MRI scans were acquired by each study site on 3T (49 sites) or 1.5T (2 sites) MRI scanners from three manufacturers (Siemens, Philips, or GE). A total of 3,290 T1w MRI scans were initially evaluated for inclusion (**Figure 1**) where participants were required to have a T1w MRI scan along with complete information on autism diagnostic status, age, and sex. Participants were excluded for ineligible diagnostic labels (autism diagnoses coded as low confidence or in remission); grossly atypical GM or WM volume estimates (GM volume <100cm^3^ or >1400cm^3^; WM volume <130cm^3^ or >800cm^3^), as based on prior work;^11^ non-independence due to family relationships (one participant per family randomly selected for inclusion); or segmentation failure (misclassified tissue), excessive motion or imaging artifacts identified through visual quality control, as in prior work.^12^ This resulted in a final analytic sample of 3,051 participants. To evaluate the robustness of the primary findings, supplementary analyses were conducted in a subsample with available FSIQ data and in two additional subsamples applying increasingly stringent motion exclusion criteria. The first motion-excluded subsample eliminated scans visually rated as having mild-moderate motion artifacts, whereas the second applied a more stringent threshold by excluding scans with any visually detectable motion artifacts.^12^

**Figure 1.**
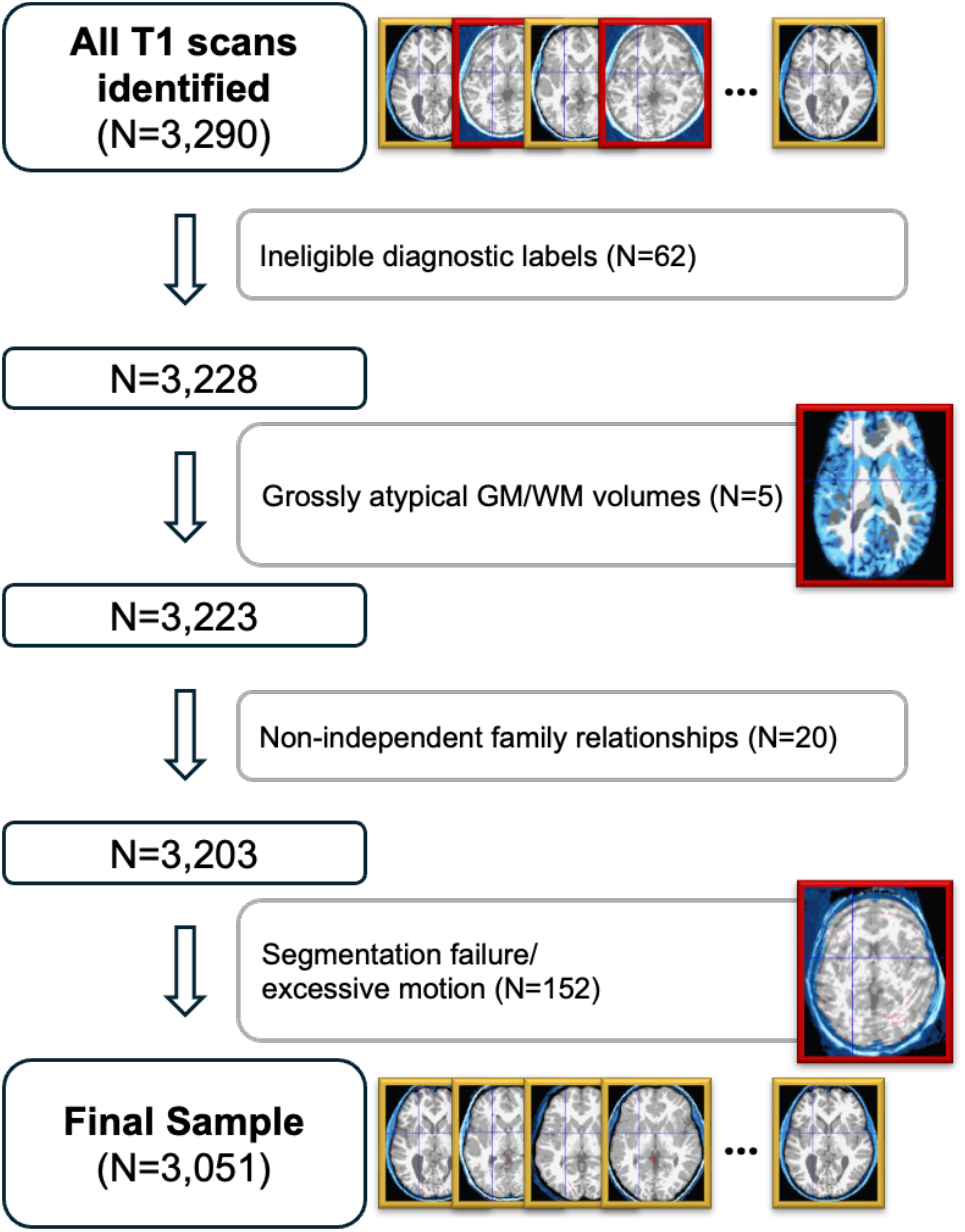
Participant selection flowchart. A total of 3,290 T1-weighted MRI scans from 51 sites/scanners were initially evaluated. Participants were excluded because of ineligible diagnostic labels (autism diagnoses coded as low confidence or in remission), grossly atypical volume estimates of gray matter (GM) or white matter (WM), non-independence of subjects due to family relationships, or segmentation failure/excessive motion, resulting in the Final Sample of 3,051 participants. Representative examples of included and excluded images are shown; images outlined in yellow passed quality control, whereas images outlined in red failed quality control and were excluded.

### T1w MRI Processing

Regional GM and WM voxel-wise volumetric morphometry were quantified using the ENIGMA CAT12 segmentation and analysis toolbox (https://neuro-jena.github.io/enigma-cat12/), as described in detail elsewhere.^12^ Overall, this pipeline uses standard Statistical Parametric Mapping (SPM) MATLAB software to perform basic preprocessing steps, followed by a large-scale voxel-wise segmentation of the 3D T1w MRI scans.^12^ Specifically, our raw scans were denoised with a spatial adaptive non-local means denoising filter, bias corrected, and affine registered. Following initial standard unified SPM segmentation, outputs were then skull-stripped and regionally parcellated. For improved local tissue intensity normalization, a Local Adaptive Segmentation intensity correction was applied, followed by an Adaptive Maximum A Posteriori segmentation, which classifies tissue type independent of tissue probability priors. Voxel classification was further refined by implementing Partial Volume Estimation, which estimates mixed-tissue fractions at tissue boundaries, thereby improving the accuracy of the resulting tissue segmentation. In the final registration step, masks were spatially normalized and modulated by the Jacobian determinants of the deformation fields, yielding the final normalized, modulated 3D whole-brain segmented GM and WM masks and volumetric estimations. These modulated GM and WM maps were spatially smoothed using a 3D Gaussian kernel (8-mm full width at half maximum, FWHM) in preparation for voxel-wise statistical analyses, consistent with previous VBM work.^11^

### Statistical Analyses

A linear mixed-effects regression analysis was conducted at each voxel across the brain to assess GM and WM volume differences between the autism and neurotypical control groups, using LONI pipeline v.7.0.3 (https://pipeline.loni.usc.edu). The model included diagnostic group as the fixed effect of interest, age, age^2^, sex, and intracranial volume (ICV) as fixed effect nuisance covariates, and site/scanner as a random effect as below:

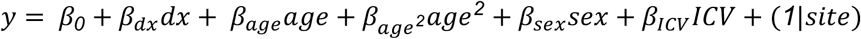

Here, *y* represents voxel-wise GM and WM volume, *dx* denotes diagnostic group (autism or neurotypical), *age* represents the linear effect of age, *age*^*2*^ is a hierarchical polynomial term for mean-centered age, and the random effect *site* represents the MRI acquisition site/scanner; for completeness, the main effects of age and sex are presented in the *Supplemental Material*. All voxel-wise *p*-values were corrected for multiple comparisons using the standard false discovery rate (FDR) with a threshold of 5% (*q*=0.05).^13^ Effect sizes are reported as standardized *β*s given the use of a linear mixed-effects model to account for nuisance covariates. To visualize the VBM results, statistical *t*-maps were thresholded at *q*=0.05, and clusters smaller than 10 voxels were excluded.^11^ Thresholded *t*-maps were then overlapped with standard atlas-based ROI masks to identify anatomical regions with significant voxels, as well as to extract regional voxel counts and peak statistics for table generation and visualization (see *Supplemental Material*).

## Results

### Participant selection and demographics

A total of 3,290 T1w MRI scans collected across 51 sites/scanners were initially identified for inclusion (**Figure 1**). As detailed in the *Methods*, participants were excluded for ineligible diagnostic labels (N=62), grossly atypical GM or WM volume estimates (N=5), non-independence of subjects due to family relationships (N=20), or segmentation failure/excessive motion (N=152), resulting in our final total sample of N=3,051 (76.8% male, 4-64 years). Demographic characteristics of the final sample consisting of 1,519 participants with autism and 1,532 neurotypical controls are presented in **Table 1**. The autism and neurotypical groups did not differ in age (*p*=0.62). The autism group had a higher proportion of males than the neurotypical group (*p*<0.001). Intracranial volume (ICV) was higher in the autism group than in the neurotypical group (*p*<0.001). Full-scale IQ (FSIQ) was lower in the autism group (*p*<0.001). To account for differences in sex distribution and brain volume between diagnosis groups, sex and ICV were included as covariates in all analyses. To assess the robustness of the findings, supplementary analyses were conducted in an FSIQ-available subsample (N=2,882), and in two additional subsamples applying increasingly stringent motion exclusion criteria, as detailed in the *Methods* (N=2,984 and N=2,815, respectively).

**Table 1.** Participant Demographics.

|  | Autism<br>(N=1,519) | Neurotypical<br>(N=1,532) | Total<br>(N=3,051) | *p*-value |
| --- | --- | --- | --- | --- |
| Age (yrs) |  |  |  |  |
| Mean (SD) | 15.1 (8.5) | 14.9 (8.0) | 15.0 (8.2) | 0.62 |
| Sex |  |  |  |  |
| Female | 262 (17.2%) | 447 (29.2%) | 709 (23.2%) | <0.001 |
| Male | 1,257 (82.8%) | 1,085 (70.8%) | 2,342 (76.8%) |  |
| FSIQ |  |  |  |  |
| Mean (SD) | 103 (18.1) | 112 (13.4) | 108 (16.5) | <0.001 |
| ICV (mL) |  |  |  |  |
| Mean (SD) | 1,540 (155) | 1,510 (141) | 1,530 (149) | <0.001 |

There were 169 (5.5%) participants missing FSIQ (72 Autism and 97 Neurotypical)

### Gray Matter Volume in Autism

The autism group exhibited lower GM volume than the neurotypical group across widespread cortical, subcortical, and cerebellar regions (**Figure 2, Figure S1, Tables S3-5)**. In the cortex, the most extensive GM volume differences were observed in the bilateral orbitofrontal cortex (OFC; t=4.92, *β*=0.08, 717 voxels), including the bilateral medial OFC, right anterior OFC, and bilateral lateral OFC, stretching into the frontal pole (t=5.25, *β*=0.08, 1,069 voxels) and subcallosal cortex (t=4.79, *β*=0.07, 221 voxels). More localized significant volume differences in autism were detected across frontal, temporal, parietal, occipital, and insular regions (**Table S3**). Among subcortical structures, the most widespread volume reductions in autism compared to the neurotypical group were in the bilateral amygdala (t=6.30, *β*=0.09, 2,278 voxels) and thalamus (t=7.39, *β*=0.13, 1,380 voxels) (**Figure 2, Tables S3-4**). Amygdala findings extended across much of both amygdalae, particularly the basolateral (t=6.09, *β*=0.08, 423 voxels) and lateral nuclei (t=6.30, *β*=0.09, 347 voxels). Within the thalamus, the mediodorsal nucleus stretching into the interthalamic adhesion (t=7.39, *β*=0.13, 790 voxels) exhibited the greatest extent of GM volume differences. In subcortical regions, smaller group difference clusters were additionally identified and listed in **Table S3-4**. Cerebellar findings were widespread, with the autism group displaying significantly lower GM volumes than the neurotypical group across many cerebellar regions (**Figure 2, Table S3, Table S5**). The largest cerebellum clusters localized to bilateral *Crus* I (t=5.33, *β*=0.09, 1,437 voxels) and *Crus* II (t=5.73, *β*=0.09, 1,787 voxels), bilateral lobules VI (t=4.76, *β*=0.07, 828 voxels), VIIb (t=4.86, *β*=0.08, 597 voxels), and VIIIa/b (t=3.91, *β*=0.08, 1516 voxels); additional cerebellar region differences are detailed in the *Supplemental Material* (**Table S3, Table S5**). Greater GM volume was observed in punctate foci within bilateral *planum polare* (t=-5.42, *β*=-0.09, 142 voxels), with additional smaller regions of greater GM volume detailed in the *Supplemental Material* (**Table S6**).

**Figure 2.**
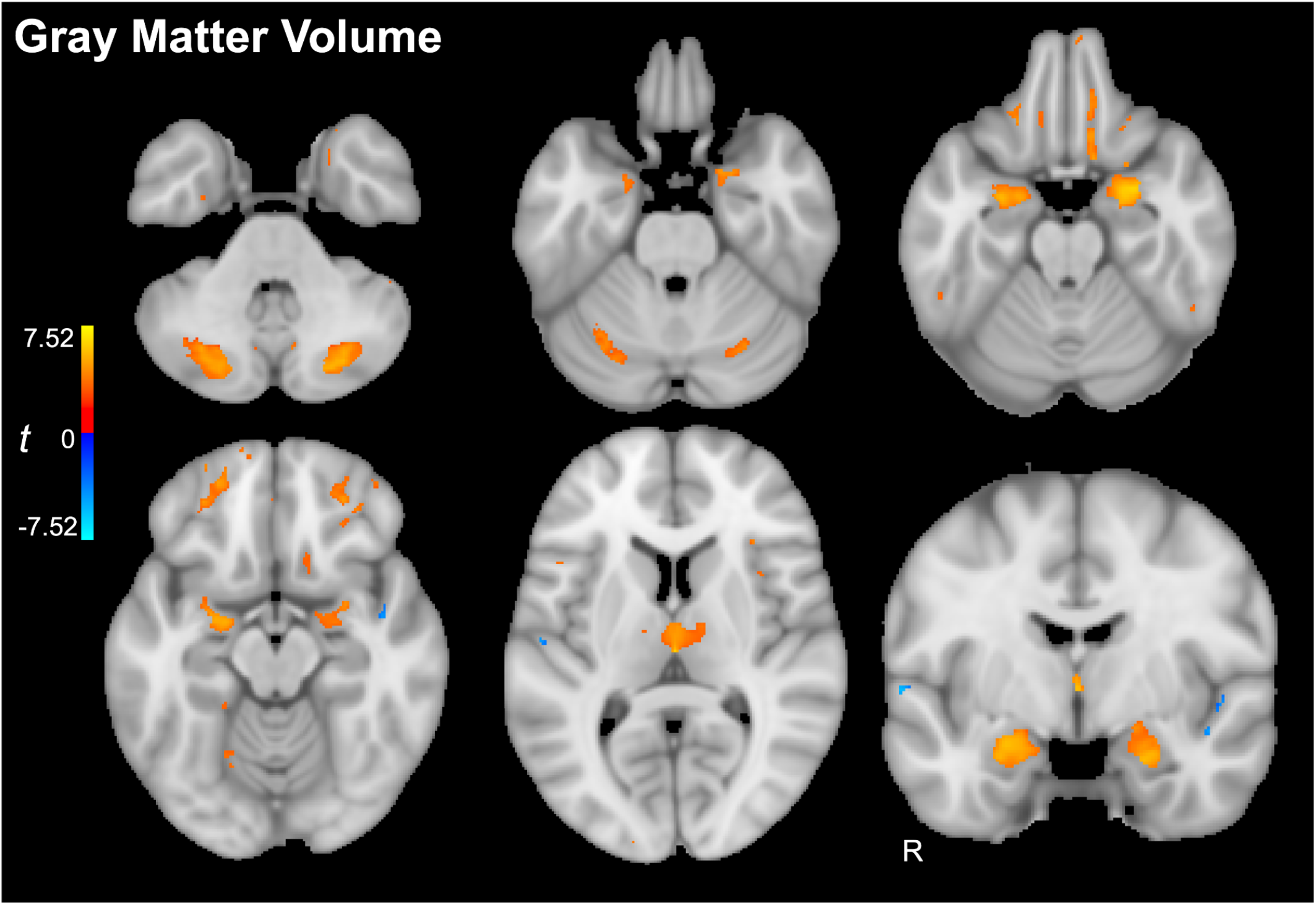
Gray matter volume differences in autism. Voxel-wise *t*-statistic map highlighting brain regions in which gray matter volume was significantly lower (red) or higher (blue) in autism than in neurotypical controls, including the orbitofrontal cortex, amygdala, thalamus, and cerebellum. See Table S3-6 for the complete list of significant regions and peak coordinates.

The overall pattern of GM findings remained largely similar following adjustment for FSIQ and application of increasingly strict motion exclusion criteria, as detailed in the *Methods* (**Figure S2-4, Table S7-18**). Across all supplementary analyses, autism continued to show widespread lower GM volumes, with the most robust differences localized to the OFC, bilateral amygdala and thalamus, and bilateral *Crus* I/II. Greater GM volume continued to be observed in a focal region of the left *planum polare*. A complete list of smaller but still significant differences is provided in the *Supplemental Material* (**Tables S7-18)**.

### White Matter Volume in Autism

The autism group displayed smaller WM volumes than the neurotypical group across widespread cerebral and cerebellar WM regions (**Figure 3, Figure S5, Table S19**). WM differences were distributed across major projection, commissural and association fibers. The most extensive differences were observed in regions that predominantly include projection fibers, particularly the bilateral anterior (t=6.74, *β*=0.08, 12,666 voxels), superior (t=5.58, *β*=0.06, 13,942 voxels), and posterior (t=4.25, *β*=0.05, 5,196 voxels) *corona radiata*. Additional regions with prominent group differences were identified in the splenium of the corpus callosum (t=5.08, *β*=0.07, 5,622 voxels) and the superior longitudinal fasciculus (t=5.51, *β*=0.06, 10,334 voxels). Affected limbic fiber regions included the cingulum (t=3.79, *β*=0.05, 342 voxels) and bilateral *fornix/stria terminalis* (t=4.57, *β*=0.07, 399 voxels). Additional significant volume differences in autism for the cortical WM are listed in **Table S19**. Significant smaller WM volumes in autism were also observed within cerebellar and brainstem WM regions, including the bilateral middle cerebellar peduncles (t=5.96, *β*=0.08, 11,553 voxels) and bilateral cortical spinal tract (t=5.61, *β*=0.07, 2651 voxels); additional cerebellar region differences are detailed in **Table S19**. One focal region of greater WM volume in autism was identified near the inferior temporal gyrus but fell outside the WM atlas and was therefore not assigned to any atlas-defined region (**Figure 3, Figure S5**)

**Figure 3.**
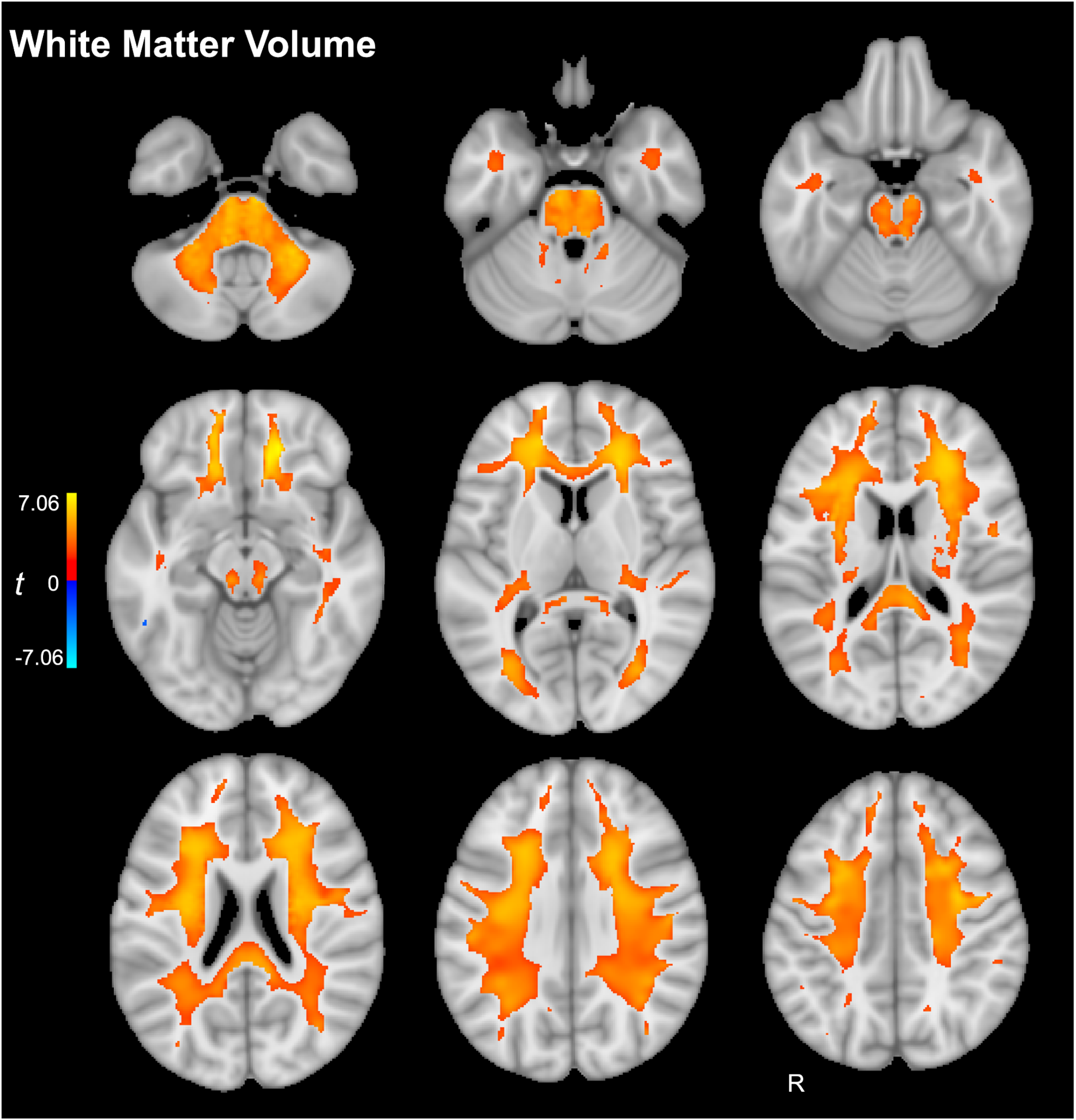
White matter volume differences in autism. Voxel-wise *t*-statistic map highlighting regions in which white matter volume was significantly lower (red) or higher (blue) in autism than in neurotypical controls, including the *corona radiata*, internal capsule, corpus callosum, and cerebellum/brainstem. See Table S19 for the complete list of significant regions and peak coordinates.

The overall pattern of WM findings remained highly consistent following adjustment for FSIQ and when applying increasingly strict motion exclusion criteria as detailed in the *Methods* (**Figure S6-8, Tables S20-22**). Across all supplementary analyses, autism continued to show smaller WM volumes across major projection, commissural, and association fiber regions as well as cerebellar WM regions; a single focal region of greater WM volume continued to be observed near the inferior temporal gyrus. A complete list of regions is provided in the *Supplemental Material* (**Tables S20-22**).

## Discussion

Here, we conducted the largest VBM mega-analysis of autism to date, leveraging structural MRI data from 3,051 participants to rigorously localize GM and WM volume differences with high spatial precision. Autism exhibited smaller GM and WM volumes across widespread cortical, subcortical, and cerebellar regions, in a pattern that remained robust after supplementary analyses. Our voxel-wise approach revealed novel GM alterations that cannot be identified using gross anatomical parcellations, such as the thalamic mediodorsal nuclei, while also precisely localizing focal GM differences within larger anatomical structures, such as the OFC. In parallel, the widespread WM alterations identified here extend previous structural MRI and diffusion MRI findings,^2,8^ suggesting convergence across complementary MRI modalities. Collectively, these findings suggest that autism is associated with structural alterations across distributed brain networks implicated in reward, sensory, and motor function.

Reward-processing theories of autism, which propose that altered reward valuation of social stimuli contributes to differences in social orienting, learning, and engagement, are supported by our observed alterations in the OFC and amygdala.^14,15^ The OFC plays a central role in reward valuation, decision-making, and social behavior, with the medial OFC supporting reward-value representation and the anterior OFC supporting identity-specific stimulus-outcome learning necessary for adaptive outcome-guided behavior.^16–18^ Through extensive connections with limbic regions, including the amygdala, the OFC contributes to evaluating the motivational significance of social information and guiding behavior in changing environments.^18^ The amygdala plays a core role in salience detection and is also involved in reward valuation, motivation, and social behavior. Accordingly, atypical amygdala development has been linked to altered social attention and social cognition in autism.^19,20^ Together, our findings support the involvement of orbitofrontal-amygdalar circuitry in autism, consistent with theories implicating reward-related neural circuitry in the condition.^14,19^

Sensory theories of autism are reinforced by our findings of alterations in the thalamus, projection fiber regions, and cerebellar regions involved in sensory processing. Sensory processing differences are prevalent in autism, with sensory theories proposing that atypical prioritization and integration of sensory and socially relevant information contribute to sensory processing differences that, in turn, affect social attention and ultimately social communication in autism.^21,22^ We identified lower GM volume localized to the interthalamic adhesion and mediodorsal thalamus, a region involved in sensory integration that also contributes to executive and affective processing through extensive reciprocal connections with cortical and limbic structures.^23^ The most extensive WM differences were observed in the *corona radiata* and internal capsule. These WM regions predominantly consist of projection fibers, including the thalamic radiations that support sensory processing and thalamo-cortical communication.^24^ Additional findings involving cerebellar lobule VI are notable given its functional connectivity with sensorimotor cortical regions.^25^ These results suggest that the atypical sensory and social processing found in autism may arise from alterations across distributed cerebello-thalamo-cortical pathways.

Motor developmental theories of autism, which propose that motor differences influence social and cognitive development, are corroborated by our present findings in motor-related cerebellar GM and WM regions.^26,27^ Motor difficulties are highly prevalent in autism and emerge early in development, with motor developmental theories proposing that atypical motor development contributes to differences in exploration and social interaction that ultimately influence social communication and behavior.^28,29^ Consistent with this framework, we observed widespread cerebellar differences involving action regions of lobule VIII, in addition to other cerebellar regions implicated in higher-order cognitive and social processes.^25,27,30^ Significant WM differences were also observed within the cerebellar peduncles, which support communication between the cerebellum, brainstem and cerebral cortex and are critical for motor control and sensorimotor integration.^31^

Collectively, our findings support network-based models of autism that emphasize alterations across reward and sensorimotor systems rather than atypical function within any single brain region. Future longitudinal studies are needed to characterize the developmental trajectories underlying the observed neuroanatomical differences in this study.^32^ Future analyses that build on the current work by using dimensional and data-driven clustering approaches may help identify biologically meaningful subgroups to further refine neurobiological models of autism.^33,34^ Complementary approaches that incorporate vertexwise measures of cortical thickness and surface area may also provide additional insights. Future studies may also benefit from refined cerebellar-specific processing pipelines to further characterize cerebellar differences associated with autism.^35^

In conclusion, this study robustly identified GM and WM differences in autism by integrating the statistical power of the largest mega-analysis of autism to date with the spatial precision of whole-brain voxel-wise analyses. Widespread alterations mapping orbitofrontal, amygdalar, thalamic, cerebellar, and major projection WM regions remained robust across multiple supplementary analyses. These findings rigorously support the involvement of distributed neural systems underlying reward processing and sensorimotor functioning in autism.

## Acknowledgments

This research was supported by the American Society of Pediatric Neuroradiology (Guerbet Research Grant Award to P.R.), National Institutes of Health (award K01MH135160 to K.E.L.) and the Asan Foundation (Biomedical Science Scholarship to G.S.K.). Research reported in this publication was also supported by the Office Of The Director, of the National Institutes of Health under Award Number S10OD032285.

Data used in the preparation of this article reside in part in the NIH-supported NIMH Data Repositories, a collaborative informatics system created by the National Institutes of Health to provide a national resource to support and accelerate research in mental health related conditions. We thank ABIDE (at http://fcon_1000.projects.nitrc.org/indi/abide) and HBN (https://doi.org/10.15387/CMI_HBN) for providing data. This manuscript reflects the views of the authors and does not reflect the opinions or views of the NIH or of the Submitters submitting original data to the NIMH Data Repositories, ABIDE or HBN.

The initial draft of the manuscript was written entirely by the authors. After the initial draft was written, ChatGPT (OpenAI) was used to assist with minor language editing of a limited number of sentences to improve readability, grammar, and phrasing only; ChatGPT was not used to generate scientific content, analyze data, or draw conclusions. The authors reviewed, edited, and verified the LLM-edited text and take full responsibility for the final text.

