## SupplementalMaterial for "Structural brain alterations in autism: A voxel-based morphometry mega-analysis in 3,051 participants across 51 sites"

### Supplemental Material


**Supplemental Methods**

**Supplemental References**

**Supplemental Figures**

- **Figure S1.** Gray matter volume in autism
- **Figure S2.** Gray matter volume in autism covarying FSIQ
- **Figure S3.** Gray matter volume in autism excluding mild-moderate motion
- **Figure S4.** Gray matter volume in autism excluding any visually detectable motion
- **Figure S5.** White matter volume in autism
- **Figure S6.** White matter volume in autism covarying FSIQ
- **Figure S7.** White matter volume in autism excluding mild-moderate motion
- **Figure S8.** White matter volume in autism excluding any visually detectable motion
- **Figure S9**. Significant main effects of age on gray and white matter
- **Figure S10**. Significant main effects of $age^2$ on gray and white matter
- **Figure S11**. Significant main effects of sex on gray and white matter

**Supplemental Tables**

- **Table S1.** Demographic information by site
- **Table S2.** Scanner information by site
- **Table S3.** Regions showing lower gray matter volumes in autism
- **Table S4.** Regions showing lower gray matter amygdalar and thalamic subnuclei volume in autism
- **Table S5.** Regions showing lower cerebellar gray matter volume in autism based on the functional parcellation
- **Table S6.** Regions showing higher gray matter volumes in autism
- **Table S7.** Regions showing lower gray matter volumes in autism covarying FSIQ
- **Table S8.** Regions showing lower gray matter amygdalar and thalamic subnuclei volume in autism covarying FSIQ
- **Table S9.** Regions showing lower cerebellar gray matter volume in autism based on the functional parcellation covarying FSIQ
- **Table S10.** Regions showing higher gray matter volumes in autism covarying FSIQ
- **Table S11.** Regions showing lower gray matter volumes in autism excluding mild-moderate motion
- **Table S12.** Regions showing lower gray matter amygdalar and thalamic subnuclei volume in autism excluding mild-moderate motion
- **Table S13.** Regions showing lower cerebellar gray matter volume in autism based on the functional parcellation excluding mild-moderate motion
- **Table S14.** Regions showing higher gray matter volumes in autism excluding mild-moderate motion
- **Table S15.** Regions showing lower gray matter volumes in autism excluding any visually detectable motion

- **Table S16.** Regions showing lower gray matter amygdalar and thalamic subnuclei volume in autism excluding any visually detectable motion
- **Table S17.** Regions showing lower cerebellar gray matter volume in autism based on the functional parcellation excluding any visually detectable motion
- **Table S18.** Regions showing higher gray matter volumes in autism excluding any visually detectable motion
- **Table S19.** Regions showing lower white matter volume in autism
- **Table S20.** Regions showing lower white matter volume in autism covarying FSIQ
- **Table S21.** Regions showing lower white matter volume in autism excluding mild-moderate motion
- **Table S22.** Regions showing lower white matter volume in autism excluding any visually detectable motion

#### Supplemental Methods

To visualize the VBM results, statistical *t*-maps were thresholded and clusters smaller than 10 voxels were excluded. Thresholded *t*-maps were then overlapped with standard atlas-based ROI masks to identify anatomical regions with significant voxels, in addition to extracting regional voxel counts and peak statistics for table generation and visualization; only atlas ROIs containing at least 10 significant voxels were included in the results tables. The atlases used include the Harvard-Oxford cortical and subcortical structural atlases for gray matter (GM);[1] the probabilistic cerebellar atlas for cerebellar GM;[2] the amygdalar subnuclei atlas by Tyszka et al.;[3] the thalamic subnuclei atlas by Najdenovska et al.;[4] the hierarchical functional cerebellar atlas;[5] and the Johns Hopkins University (JHU) atlas for white matter (WM).[6]

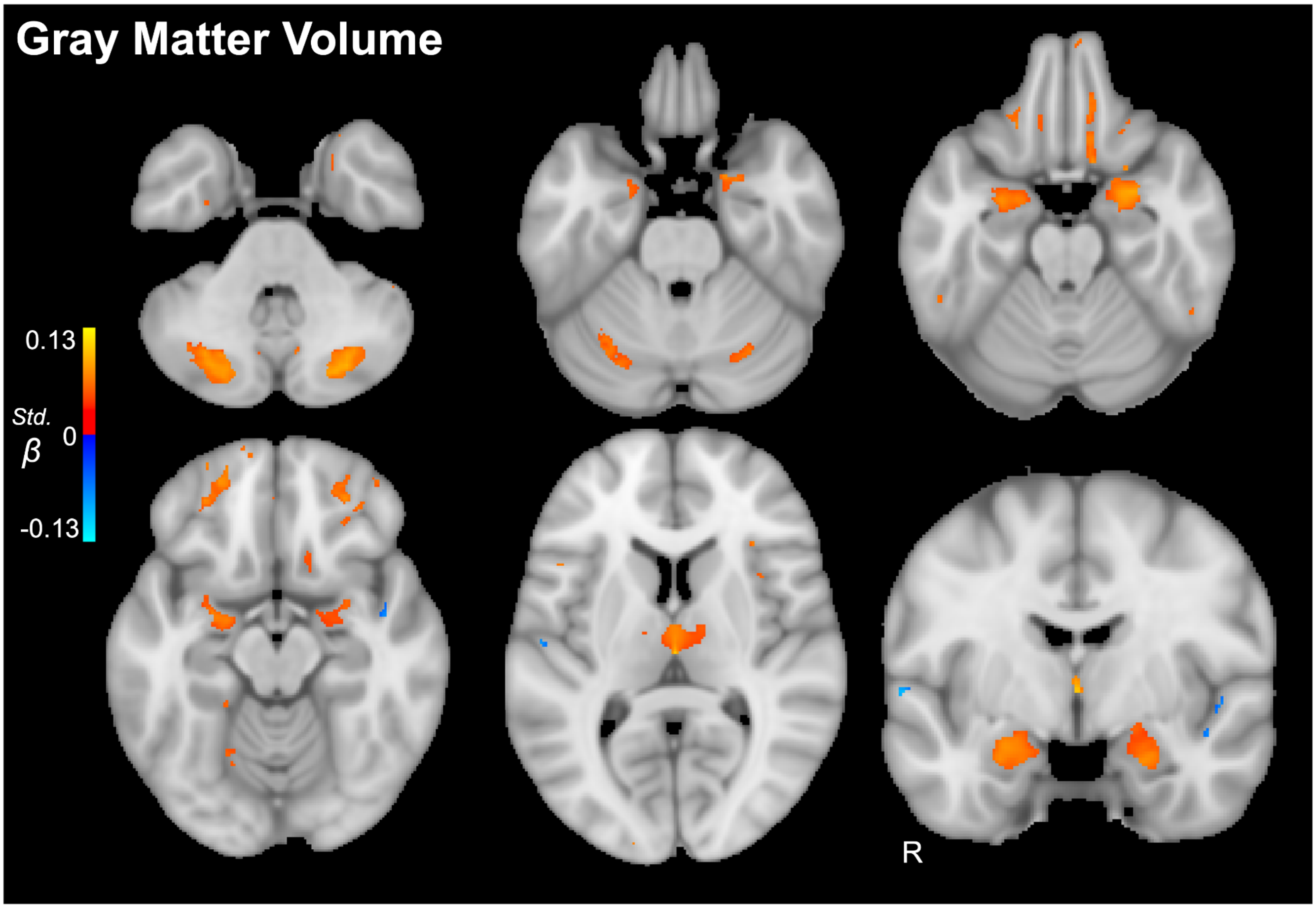


**Figure S1. Gray matter volume in autism**

Voxel-wise standardized *β*-statistic map highlighting brain regions in which gray matter volume was significantly lower (red) or higher (blue) in autism than in neurotypical controls. The significant regions included the orbitofrontal cortex, amygdala, thalamus, and cerebellum. See Table S3-6 for the complete list of significant regions and peak coordinates.

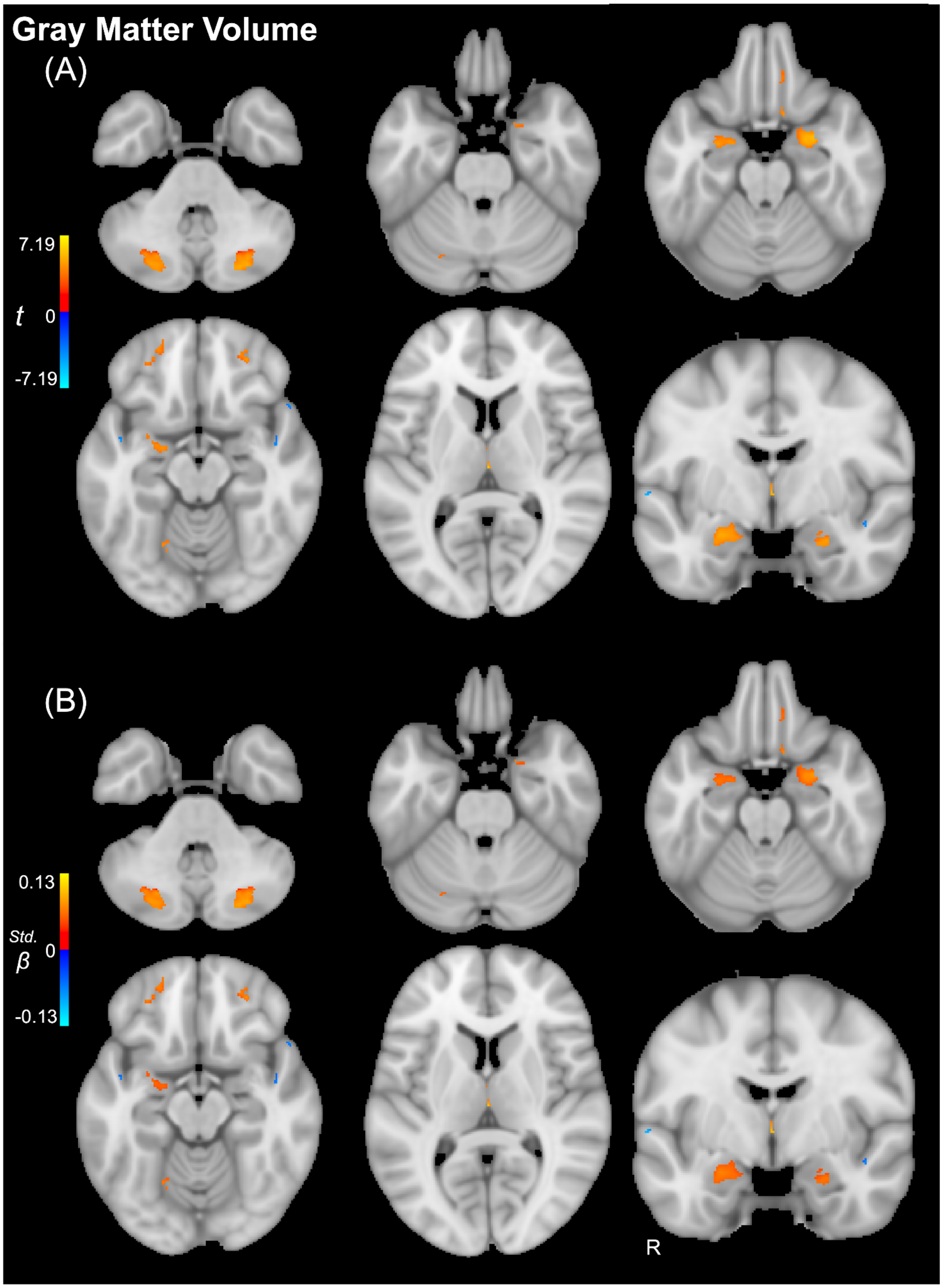


**Figure S2. Gray matter volume in autism covarying FSIQ**

Voxel-wise (A) *t*-statistic map and (B) standardized *β*-statistic map highlighting brain regions in which gray matter volume was significantly lower (red) or higher (blue) in autism than in neurotypical controls covarying full-scale IQ. The significant regions included the orbitofrontal cortex, amygdala, thalamus, and cerebellum. See Table S7-10 for the complete list of significant regions and peak coordinates.

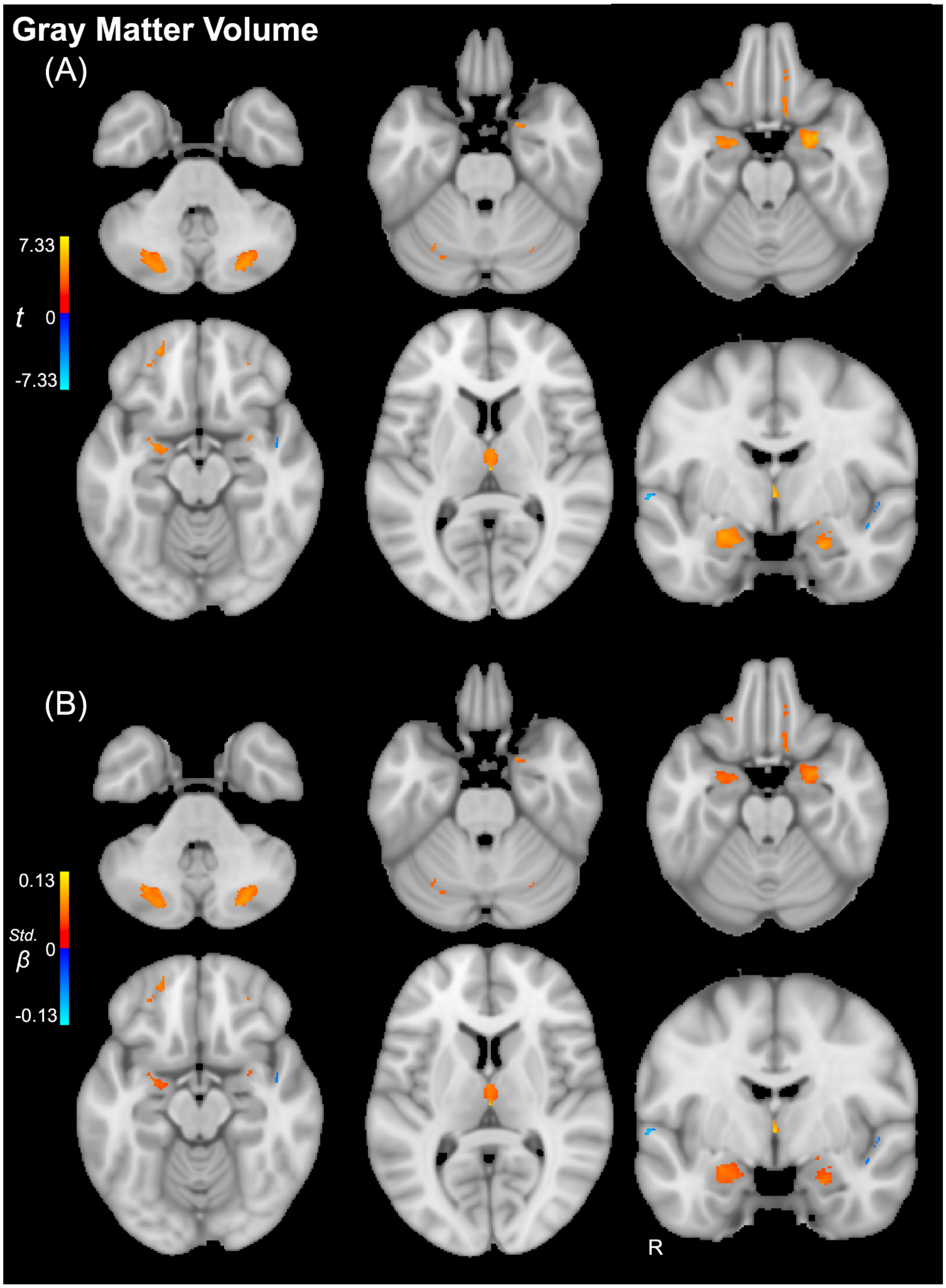


**Figure S3. Gray matter volume in autism excluding mild-moderate motion**

Voxel-wise (A) *t*-statistic map and (B) standardized *β*-statistic map highlighting brain regions in which gray matter volume was significantly lower (red) or higher (blue) in autism than in neurotypical controls after excluding participants with mild-moderate motion. The significant regions included the orbitofrontal cortex, amygdala, thalamus, and cerebellum. See Table S11-14 for the complete list of significant regions and peak coordinates.

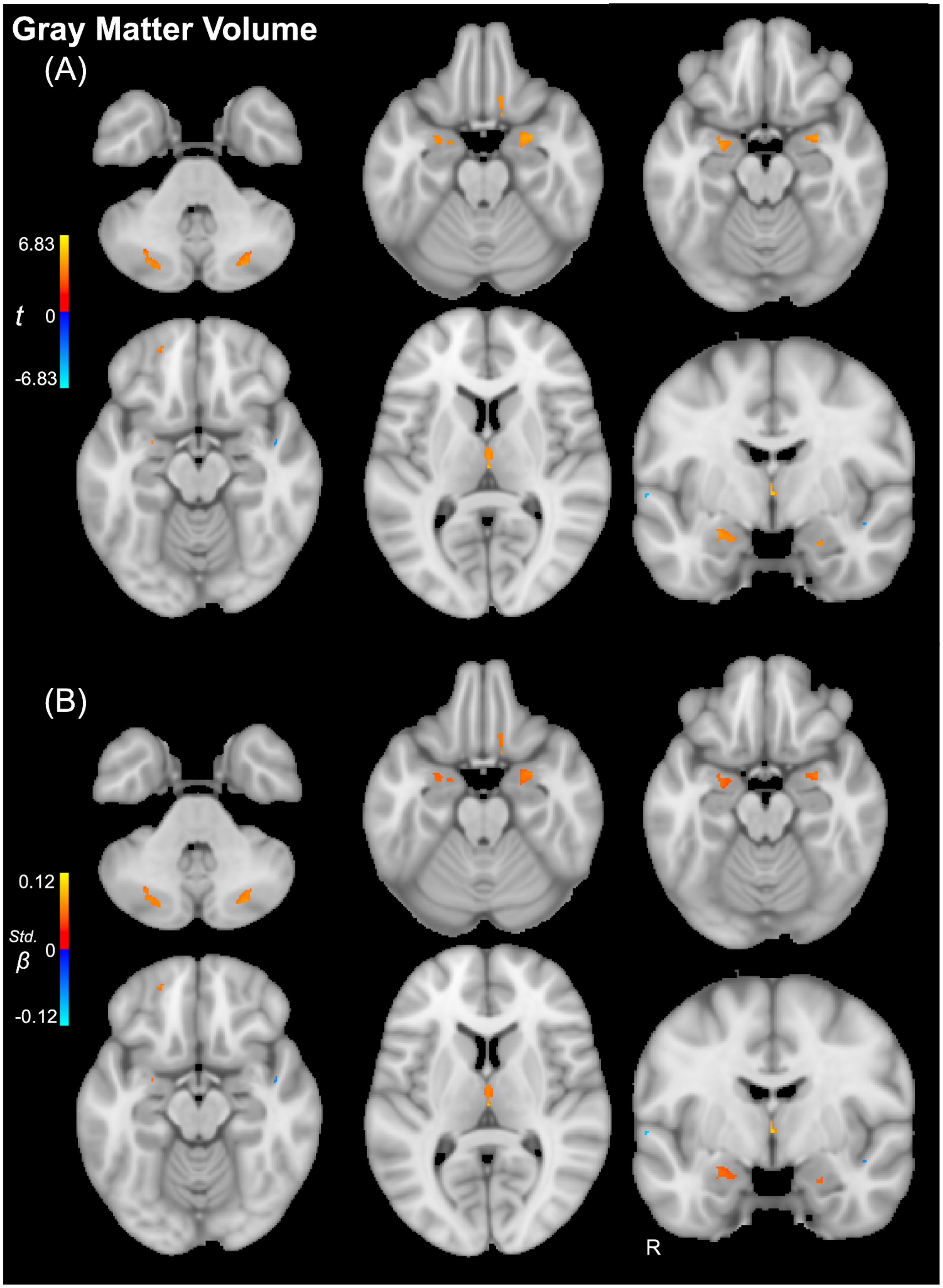


**Figure S4. Gray matter volume in autism excluding any visually detectable motion**

Voxel-wise (A) *t*-statistic map and (B) standardized *β*-statistic map highlighting brain regions in which gray matter volume was significantly lower (red) or higher (blue) in autism than in neurotypical controls after excluding participants with any visually detectable motion. The significant regions included the orbitofrontal cortex, amygdala, thalamus, and cerebellum. See Table S15-18 for the complete list of significant regions and peak coordinates.

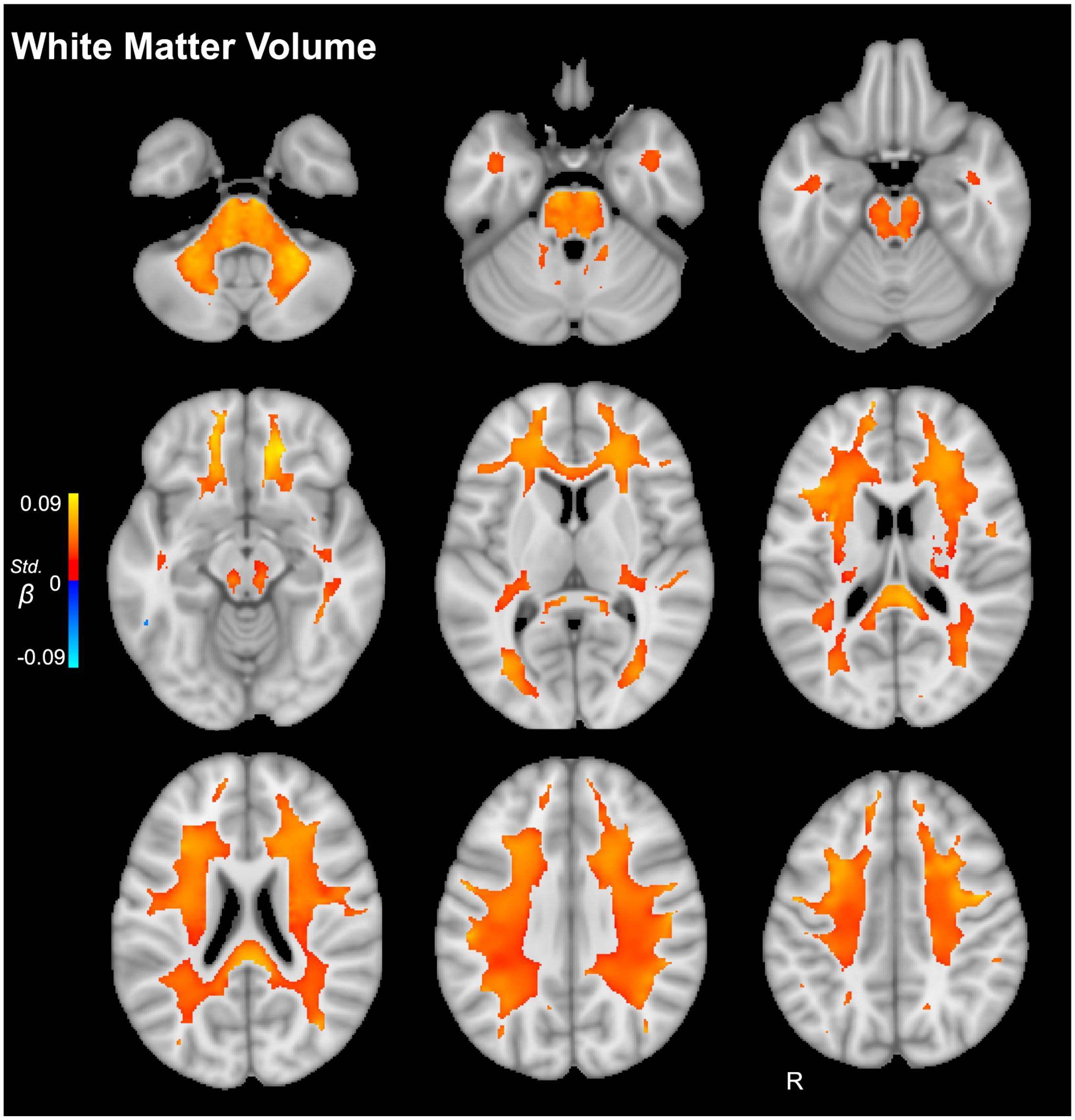


**Figure S5. White matter volume in autism**

Voxel-wise standardized $\beta$-statistic map highlighting regions in which white matter volume was significantly lower (red) or higher (blue) in autism than in neurotypical controls. The significant regions included *corona radiata,* internal capsule, corpus callosum, and cerebellum/brainstem. See Table S19 for the complete list of significant regions and peak coordinates.

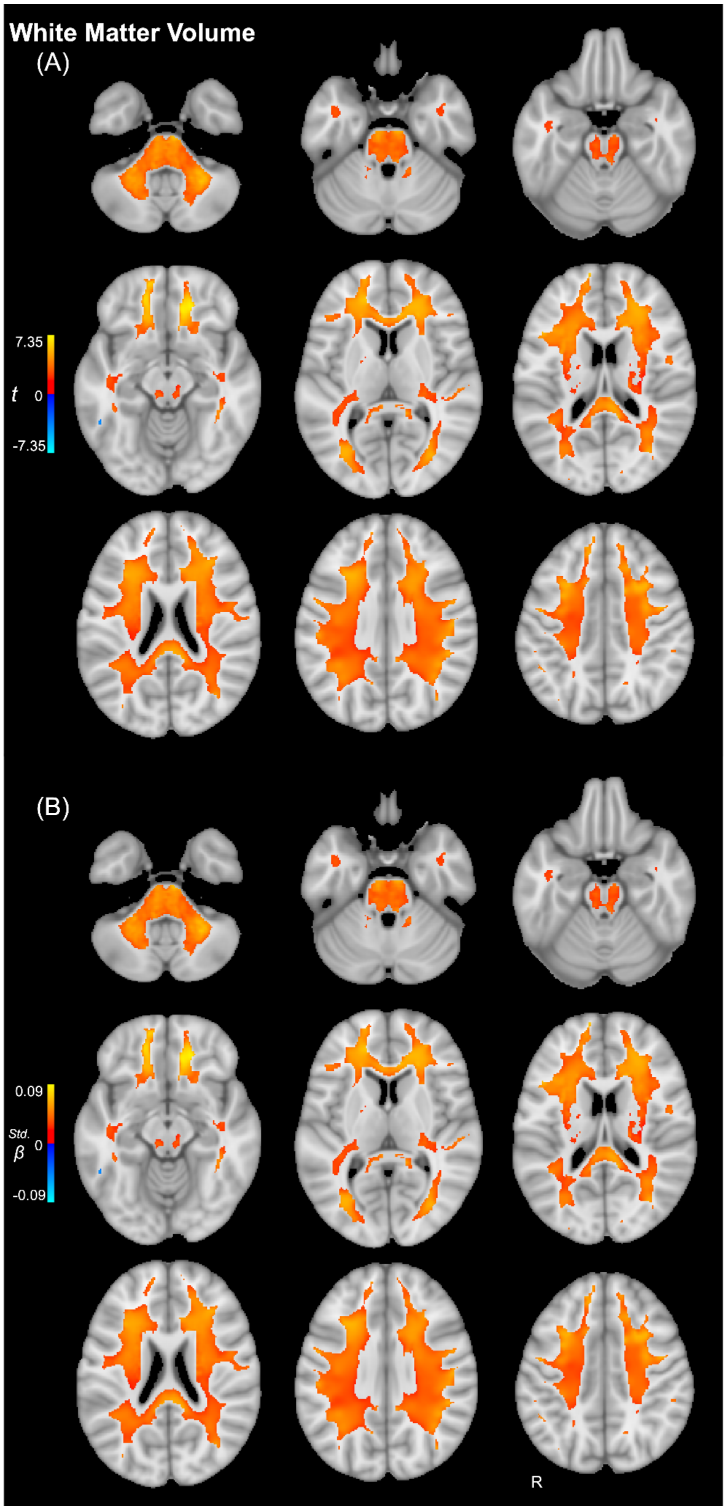


**Figure S6. White matter volume in autism covarying FSIQ**

Voxel-wise (A) *t*-statistic map and (B) standardized *β*-statistic map highlighting regions in which white matter volume was significantly lower (red) or higher (blue) in autism than in neurotypical controls covarying full-scale IQ. The significant regions included *corona radiata,* internal capsule, corpus callosum, and cerebellum/brainstem. See Table S20 for the complete list of significant regions and peak coordinates.

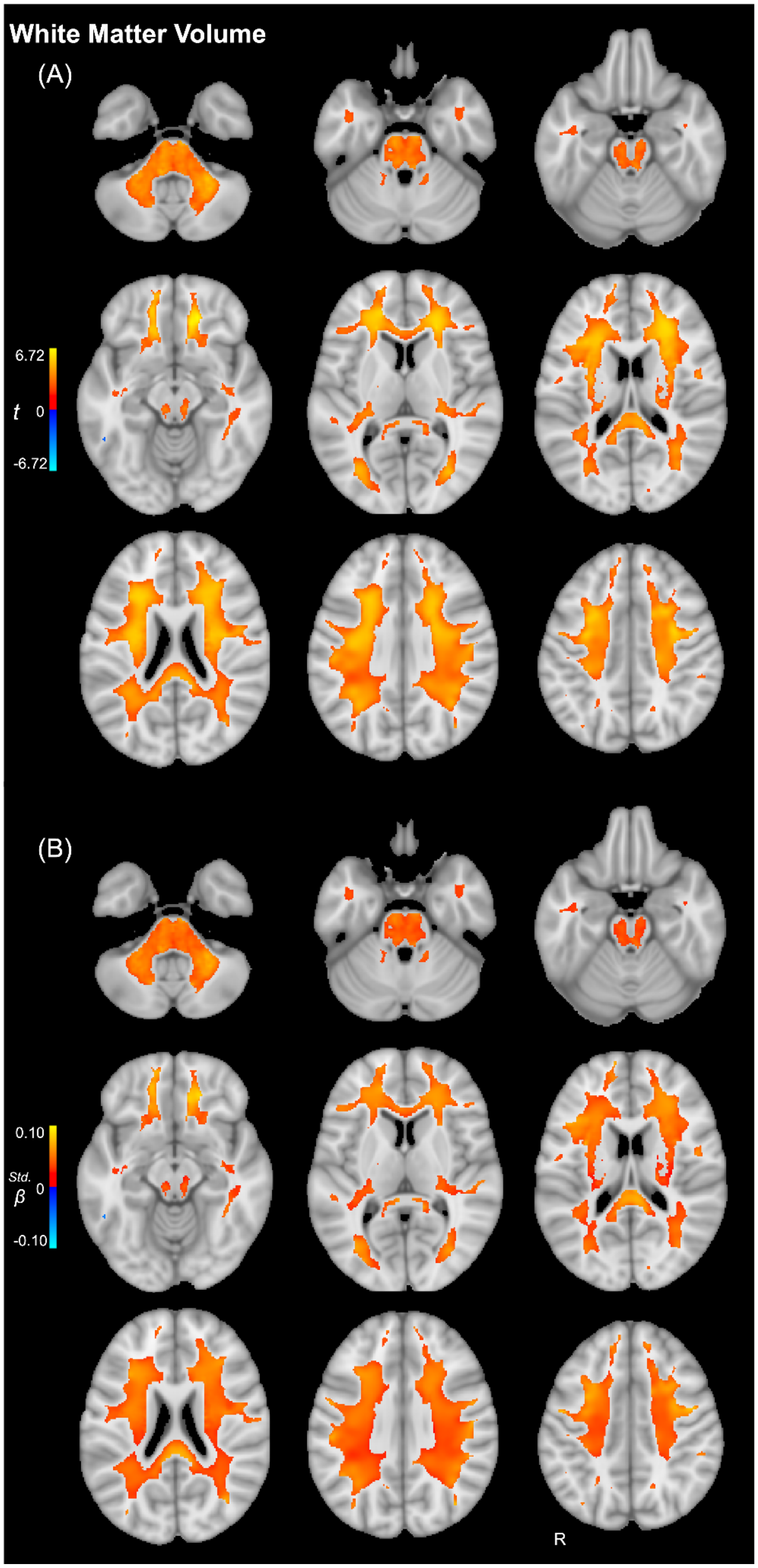


**Figure S7. White matter volume in autism excluding mild-moderate motion**

Voxel-wise (A) *t*-statistic map and (B) standardized *β*-statistic map highlighting regions in which white matter volume was significantly lower (red) or higher (blue) in autism than in neurotypical controls after excluding participants with mild-moderate motion. The significant regions included *corona radiata,* internal capsule, corpus callosum, and cerebellum/brainstem. See Table S21 for the complete list of significant regions and peak coordinates.

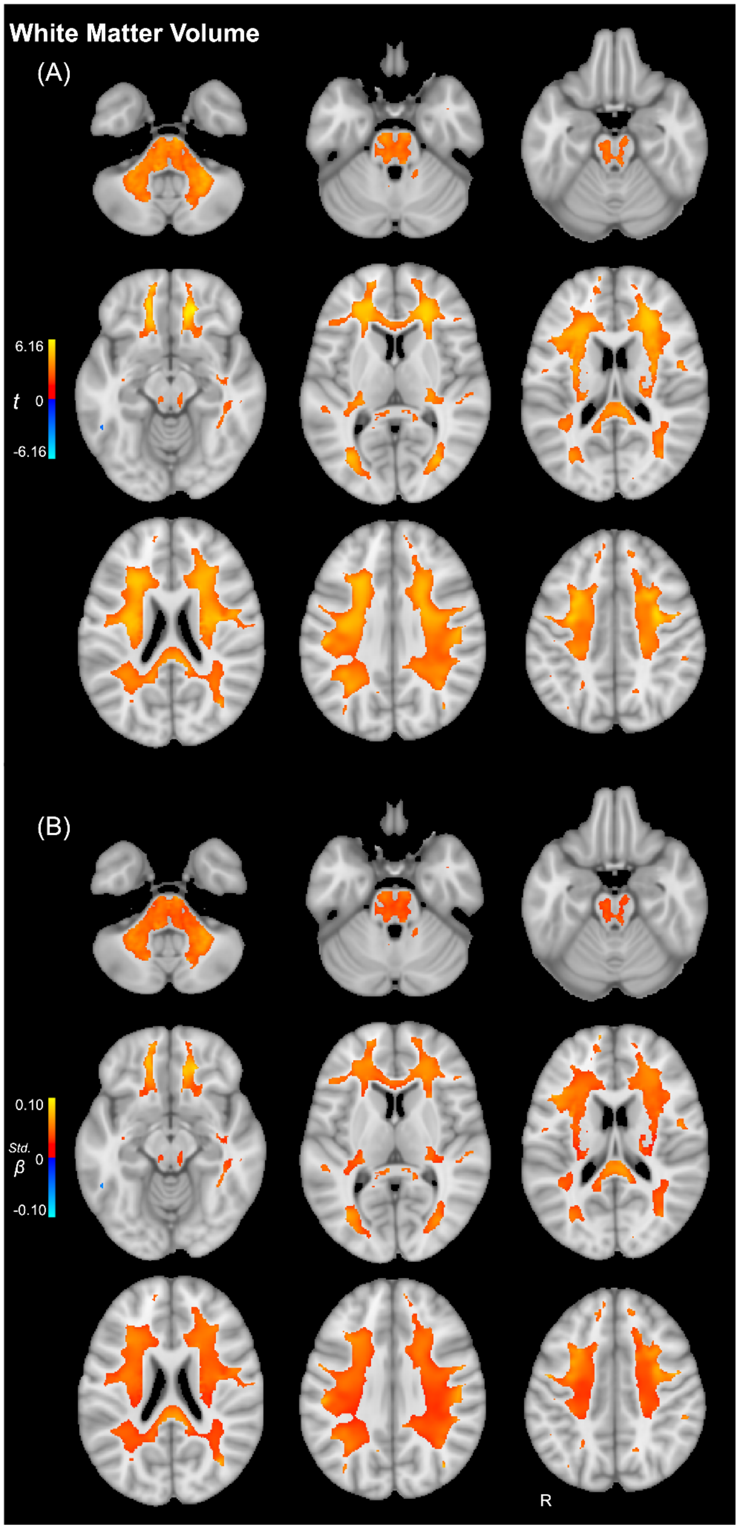


**Figure S8. White matter volume in autism excluding any visually detectable motion**

Voxel-wise (A) *t*-statistic map and (B) standardized *β*-statistic map highlighting regions in which white matter volume was significantly lower (red) or higher (blue) in autism than in neurotypical controls after excluding participants with any visually detectable motion. The significant regions included *corona radiata,* internal capsule, corpus callosum, and cerebellum/brainstem. See Table S22 for the complete list of significant regions and peak coordinates.

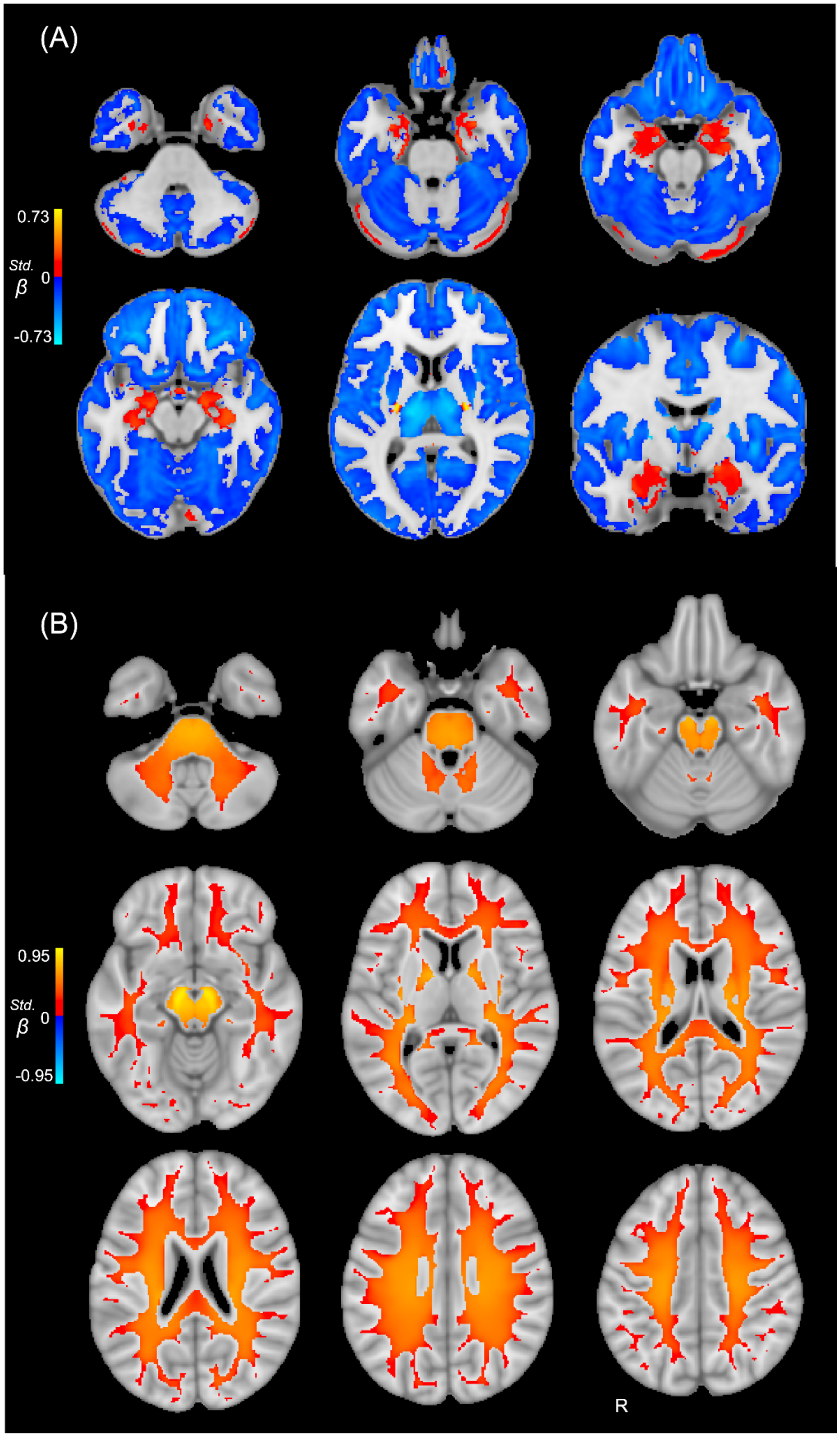


**Figure S9. Significant main effects of age on gray and white matter**

Voxel-wise standardized $\beta$-statistic map highlighting (A) gray and (B) white matter regions showing significant linear age effects after accounting for the quadratic age term. Linear and quadratic age effects should be interpreted jointly. This figure is shown for completeness and the primary analyses focused on diagnosis differences shown in Figures 2-3.

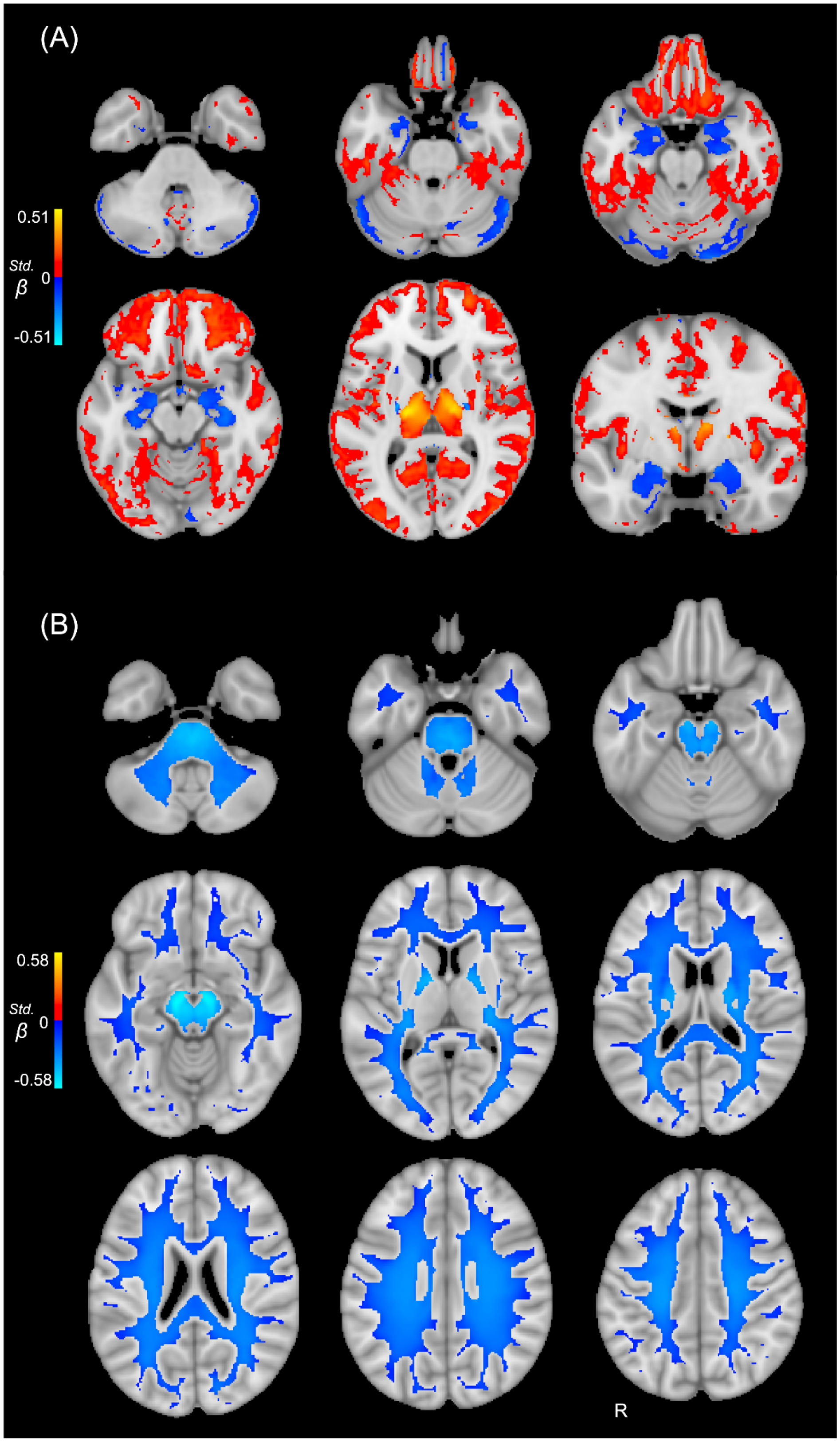


**Figure S10. Significant main effects of age$^2$ on gray and white matter**

Voxel-wise standardized $\beta$-statistic map highlighting (A) gray and (B) white matter regions showing significant quadratic age effects (age$^2$). Linear and quadratic age effects should be interpreted jointly. This figure is shown for completeness and the primary analyses focused on diagnosis differences shown in Figures 2-3.

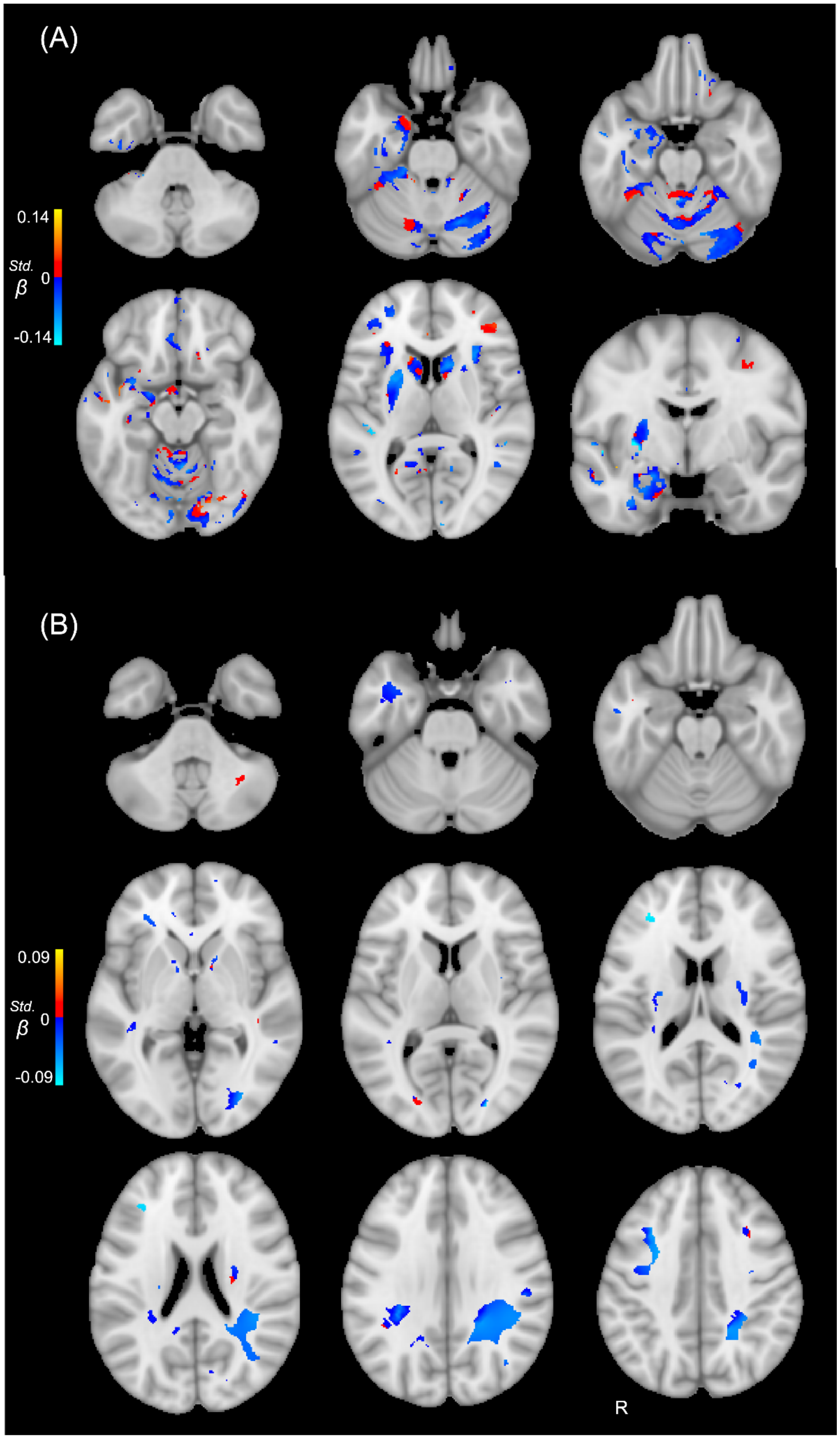


**Figure S11. Significant main effects of sex on gray and white matter**

Voxel-wise standardized $\beta$-statistic map highlighting (A) gray and (B) white matter regions showing significant associations with sex. Red indicates greater volume in females than males, whereas blue indicates greater values in males than females. This figure is shown for completeness and the primary analyses focused on diagnosis differences shown in Figures 2-3.

**Table S1. Demographic information by site**

| Site/Scanner | N | Autism | Neurotypical | Female | Male | Age (years) | ICV ($cm^3$) | Full-Scale IQ[a] | ADOS[a] CSS | SRS[a] T-Score |
|---|---|---|---|---|---|---|---|---|---|---|
| ABIDEI-CALTECH | 38 | 19 (50.0%) | 19 (50.0%) | 8 (21.1%) | 30 (78.9%) | 28.2±10.6 | 1581.0±165.9 | 111.3±11.4 | NA | NA |
| ABIDEI-CMU | 27 | 14 (51.9%) | 13 (48.1%) | 6 (22.2%) | 21 (77.8%) | 26.6±5.7 | 1564.2±174.3 | 114.6±10.5 | NA | NA |
| ABIDEI-KKI | 54 | 22 (40.7%) | 32 (59.3%) | 13 (24.1%) | 41 (75.9%) | 10.1±1.3 | 1598.7±142.4 | 106.9±15.4 | 8.1±1.8 | NA |
| ABIDEI-Leuven_1 | 29 | 14 (48.3%) | 15 (51.7%) | 0 (0.0%) | 29 (100.0%) | 22.6±3.6 | 1588.1±112.0 | 112.2±13.0 | NA | 56.8±9.6 |
| ABIDEI-Leuven_2 | 35 | 15 (42.9%) | 20 (57.1%) | 8 (22.9%) | 27 (77.1%) | 14.2±1.4 | 1519.9±109.0 | NA | NA | 56.7±17.3 |
| ABIDEI-MaxMun | 57 | 24 (42.1%) | 33 (57.9%) | 7 (12.3%) | 50 (87.7%) | 26.2±12.1 | 1548.1±123.9 | 110.8±11.3 | NA | NA |
| ABIDEI-NYU | 184 | 79 (42.9%) | 105 (57.1%) | 37 (20.1%) | 147 (79.9%) | 15.3±6.6 | 1499.5±151.9 | 110.9±14.9 | 7.0±2.1 | 58.9±16.0 |
| ABIDEI-OHSU | 28 | 13 (46.4%) | 15 (53.6%) | 0 (0.0%) | 28 (100.0%) | 10.8±1.9 | 1536.3±125.3 | 111.6±16.9 | NA | NA |
| ABIDEI-OLIN | 33 | 17 (51.5%) | 16 (48.5%) | 5 (15.2%) | 28 (84.8%) | 16.8±3.4 | 1537.8±155.5 | 113.2±17.5 | 9.0±0.0 | NA |
| ABIDEI-PITT | 57 | 30 (52.6%) | 27 (47.4%) | 8 (14.0%) | 49 (86.0%) | 18.9±6.9 | 1523.4±156.3 | 110.1±12.2 | NA | NA |
| ABIDEI-SBL | 29 | 15 (51.7%) | 14 (48.3%) | 0 (0.0%) | 29 (100.0%) | 34.6±8.6 | 1561.3±83.0 | 109.2±13.6 | NA | NA |
| ABIDEI-SDSU | 36 | 14 (38.9%) | 22 (61.1%) | 7 (19.4%) | 29 (80.6%) | 14.4±1.8 | 1579.9±145.5 | 109.4±13.8 | 6.5±1.9 | NA |
| ABIDEI-Stanford | 40 | 20 (50.0%) | 20 (50.0%) | 8 (20.0%) | 32 (80.0%) | 10.0±1.6 | 1501.6±131.6 | 112.3±16.4 | 7.5±1.9 | NA |
| ABIDEI-Trinity | 46 | 24 (52.2%) | 22 (47.8%) | 0 (0.0%) | 46 (100.0%) | 17.0±3.5 | 1609.3±136.1 | 110.0±13.7 | NA | NA |
| ABIDEI-UCLA_1 | 70 | 40 (57.1%) | 30 (42.9%) | 10 (14.3%) | 60 (85.7%) | 13.2±2.4 | 1552.9±131.1 | 103.7±12.1 | 6.3±2.0 | NA |
| ABIDEI-UCLA_2 | 24 | 12 (50.0%) | 12 (50.0%) | 2 (8.3%) | 22 (91.7%) | 12.6±1.5 | 1598.0±128.6 | 102.4±14.3 | 8.1±1.3 | NA |
| ABIDEI-UM_1 | 105 | 50 (47.6%) | 55 (52.4%) | 25 (23.8%) | 80 (76.2%) | 13.5±2.9 | 1501.2±145.6 | 105.6±14.2 | 6.5±2.2 | NA |
| ABIDEI-UM_2 | 35 | 13 (37.1%) | 22 (62.9%) | 2 (5.7%) | 33 (94.3%) | 16.0±3.3 | 1485.5±144.9 | 112.2±10.7 | 6.5±2.3 | NA |
| ABIDEI-USM | 96 | 55 (57.3%) | 41 (42.7%) | 0 (0.0%) | 96 (100.0%) | 22.1±7.8 | 1603.2±130.4 | 106.5±16.7 | 5.3±4.0 | 58.8±17.4 |
| ABIDEI-Yale | 55 | 27 (49.1%) | 28 (50.9%) | 15 (27.3%) | 40 (72.7%) | 12.6±2.9 | 1486.4±153.0 | 100.1±20.2 | 7.1±1.7 | 60.2±16.7 |

| Site/Scanner | N | Autism | Neurotypical | Female | Male | Age (years) | ICV ($cm^3$) | Full-Scale IQ[a] | ADOS[a] CSS | SRS[a] T-Score |
|---|---|---|---|---|---|---|---|---|---|---|
| ABIDEII-BNI_1 | 56 | 29 (51.8%) | 27 (48.2%) | 0 (0.0%) | 56 (100.0%) | 38.3±15.7 | 1528.9±117.1 | 109.8±13.1 | NA | 59.7±16.3 |
| ABIDEII-EMC_1 | 51 | 24 (47.1%) | 27 (52.9%) | 10 (19.6%) | 41 (80.4%) | 8.2±1.1 | 1505.1±156.1 | NA | NA | NA |
| ABIDEII-ETH_1 | 33 | 11 (33.3%) | 22 (66.7%) | 0 (0.0%) | 33 (100.0%) | 22.7±4.5 | 1565.7±114.2 | 113.1±11.5 | NA | 54.3±13.9 |
| ABIDEII-GU_1 | 101 | 49 (48.5%) | 52 (51.5%) | 34 (33.7%) | 67 (66.3%) | 10.7±1.6 | 1486.8±137.9 | 120.6±14.1 | 6.1±2.5 | 58.5±17.9 |
| ABIDEII-IP_1 | 54 | 22 (40.7%) | 32 (59.3%) | 29 (53.7%) | 25 (46.3%) | 20.5±10.4 | 1468.5±131.5 | 95.0±23.4 | 8.3±1.5 | NA |
| ABIDEII-IU_1 | 40 | 20 (50.0%) | 20 (50.0%) | 9 (22.5%) | 31 (77.5%) | 24.4±7.4 | 1604.0±156.2 | 116.7±11.1 | NA | 62.3±13.6 |
| ABIDEII-KKI_1 | 210 | 55 (26.2%) | 155 (73.8%) | 70 (33.3%) | 140 (66.7%) | 10.3±1.3 | 1559.1±149.0 | 111.5±13.1 | 6.9±1.8 | 51.8±14.9 |
| ABIDEII-KUL_3 | 21 | 21 (100.0%) | 0 (0.0%) | 0 (0.0%) | 21 (100.0%) | 22.5±3.4 | 1560.7±144.9 | 108.1±15.2 | NA | 58.1±7.9 |
| ABIDEII-NYU_1 | 76 | 46 (60.5%) | 30 (39.5%) | 7 (9.2%) | 69 (90.8%) | 9.9±5.0 | 1494.8±108.4 | 107.7±17.9 | 6.3±2.1 | 62.3±17.5 |
| ABIDEII-OHSU_1 | 93 | 37 (39.8%) | 56 (60.2%) | 36 (38.7%) | 57 (61.3%) | 10.9±2.0 | 1504.6±151.2 | 112.9±15.1 | 6.8±1.9 | 58.6±17.1 |
| ABIDEII-ONRC_2 | 57 | 22 (38.6%) | 35 (61.4%) | 18 (31.6%) | 39 (68.4%) | 23.2±3.8 | 1535.2±160.1 | 112.0±14.2 | NA | NA |
| ABIDEII-SDSU_1 | 56 | 32 (57.1%) | 24 (42.9%) | 9 (16.1%) | 47 (83.9%) | 13.2±3.1 | 1580.1±143.2 | 101.1±13.6 | 7.9±1.8 | 64.1±19.3 |
| ABIDEII-STANFORD | 38 | 19 (50.0%) | 19 (50.0%) | 3 (7.9%) | 35 (92.1%) | 11.1±1.2 | 1637.9±145.0 | 113.3±14.8 | 6.9±1.6 | NA |
| ABIDEII-TCD_1 | 42 | 21 (50.0%) | 21 (50.0%) | 0 (0.0%) | 42 (100.0%) | 15.2±3.2 | 1577.4±125.2 | 113.5±15.0 | NA | 60.1±17.2 |
| ABIDEII-UCD_1 | 31 | 17 (54.8%) | 14 (45.2%) | 7 (22.6%) | 24 (77.4%) | 14.8±1.9 | 1610.3±138.5 | 107.9±12.6 | 6.2±1.8 | 58.4±18.0 |
| ABIDEII-UCLA_1 | 23 | 12 (52.2%) | 11 (47.8%) | 5 (21.7%) | 18 (78.3%) | 10.9±2.3 | 1527.9±113.4 | 107.9±15.7 | 7.8±1.9 | NA |
| ABIDEII-UM | 25 | 11 (44.0%) | 14 (56.0%) | 6 (24.0%) | 19 (76.0%) | 10.0±2.0 | 1459.0±144.2 | 112.5±16.8 | 6.6±2.2 | NA |
| ABIDEII-USM_1 | 30 | 15 (50.0%) | 15 (50.0%) | 5 (16.7%) | 25 (83.3%) | 21.8±7.7 | 1547.6±153.4 | 109.0±19.3 | NA | 62.0±19.2 |
| HBN-CBIC | 200 | 135 (67.5%) | 65 (32.5%) | 48 (24.0%) | 152 (76.0%) | 10.7±3.7 | 1502.1±152.0 | 101.6±18.0 | NA | 61.2±13.1 |
| HBN-CUNY | 59 | 39 (66.1%) | 20 (33.9%) | 11 (18.6%) | 48 (81.4%) | 10.7±3.4 | 1502.4±155.6 | 103.1±17.2 | NA | 61.0±13.4 |
| HBN-RU | 199 | 98 (49.2%) | 101 (50.8%) | 56 (28.1%) | 143 (71.9%) | 10.4±3.7 | 1452.3±138.7 | 99.8±17.2 | NA | 60.3±14.5 |

| Site/Scanner | N | Autism | Neurotypical | Female | Male | Age (years) | ICV ($cm^3$) | Full-Scale IQ[a] | ADOS[a] CSS | SRS[a] T-Score |
|---|---|---|---|---|---|---|---|---|---|---|
| HBN-SI | 87 | 34 (39.1%) | 53 (60.9%) | 35 (40.2%) | 52 (59.8%) | 11.9±3.5 | 1492.6±143.5 | 101.7±17.0 | NA | 57.9±15.2 |
| NDA1906 | 70 | 54 (77.1%) | 16 (22.9%) | 1 (1.4%) | 69 (98.6%) | 18.6±9.7 | 1597.2±148.2 | 101.1±17.0 | NA | 68.6±21.2 |
| NDA2021_HT | 52 | 31 (59.6%) | 21 (40.4%) | 20 (38.5%) | 32 (61.5%) | 12.3±3.1 | 1513.8±160.8 | 104.8±18.0 | NA | 61.0±16.2 |
| NDA2021_SP | 57 | 25 (43.9%) | 32 (56.1%) | 25 (43.9%) | 32 (56.1%) | 13.0±2.7 | 1503.8±156.5 | 110.2±21.6 | NA | 57.4±18.1 |
| NDA2021_ST | 28 | 18 (64.3%) | 10 (35.7%) | 16 (57.1%) | 12 (42.9%) | 11.9±2.9 | 1505.7±139.6 | 112.2±18.7 | NA | 64.2±18.0 |
| NDA2021_UP | 26 | 10 (38.5%) | 16 (61.5%) | 14 (53.8%) | 12 (46.2%) | 12.6±3.3 | 1525.0±166.9 | 106.6±16.9 | NA | 56.3±18.7 |
| NDA2021_UT | 69 | 40 (58.0%) | 29 (42.0%) | 30 (43.5%) | 39 (56.5%) | 12.9±3.1 | 1485.6±136.7 | 103.8±17.4 | NA | 61.6±16.3 |
| NDA2021_YT | 65 | 32 (49.2%) | 33 (50.8%) | 40 (61.5%) | 25 (38.5%) | 13.5±2.8 | 1475.8±139.5 | 104.9±17.2 | NA | 59.6±16.3 |
| NDA2291 | 13 | 13 (100.0%) | 0 (0.0%) | 3 (23.1%) | 10 (76.9%) | 48.5±6.8 | 1485.9±141.3 | 104.7±21.4 | NA | NA |
| NDA2834 | 11 | 10 (90.9%) | 1 (9.1%) | 1 (9.1%) | 10 (90.9%) | 9.2±1.3 | 1524.6±152.3 | 103.5±30.9 | NA | 76.5±5.2 |

FSIQ, Full Scale Intelligence Quotient; ADOS CSS, Autism Diagnostic Observation Schedule Calibrated Severity Score; SRS T-score, Social Responsiveness Scale T-score. [a]The mean value is calculated based only on subjects with available data for this measure.

**Table S2. Scanner information by site**

| Site/Scanner | Vendor | Scanner | Field strength (T) | Voxel Size (mm) | TE (ms) | TR (ms) | FOV (mm) |
|---|---|---|---|---|---|---|---|
| NDA1906 | Siemens | Trio | 3 | 1.0x1.0x1.0 | 2.93 | 1800 | 256 |
| NDA2021-HT | Siemens | TrioTim | 3 | 1.0x1.0x1.0 | 3.31 | 2530 | 256 |
| NDA2021-SP | Siemens | Prisma | 3 | 1.0x1.0x1.0 | 3.34 | 2530 | 256 |
| NDA2021-ST | Siemens | TrioTim | 3 | 1.0x1.0x1.0 | 3.31 | 2530 | 256 |
| NDA2021-UP | Siemens | Prisma | 3 | 1.0x1.0x1.0 | 3.34 | 2530 | 256 |
| NDA2021-UT | Siemens | TrioTim | 3 | 1.0x1.0x1.0 | 3.31 | 2530 | 256 |
| NDA2021-YT | Siemens | TrioTim | 3 | 1.0x1.0x1.0 | 3.31 | 2530 | 256 |
| NDAR 2291 | GE | MR750 | 3 | 0.5x0.5x0.8 | 3.66 | 8.78 | 256 |
| NDAR 2834 | Siemens | Allegra | 3 | 0.8x0.8x0.8 | 4.35 | 2500 | 256 |
| ABIDEI-CALTECH | Siemens | MAGNETOM TrioTim | 3T | 1.0x1.0x1.0 | 2.73 | 1590 | 256 |
| ABIDEI-CMU | Siemens | MAGNETOM Verio | 3T | 1.0x1.0x1.0 | 2.48 | 1870 | 256 |
| ABIDEI-KKI | Philips | Achieva | 3T | 1.0x1.0x1.0 | 3.7 | 8.0 | 256 |
| ABIDEI-Leuven1 | Philips | Interna | 3T | .98x.98x1.2 | 3.6 | 9.6 | 250 |
| ABIDEI-Leuven2 | Philips | Interna | 3T | .98x.98x1.2 | 4.6 | 9.6 | 250 |
| ABIDEI-MaxMun | Siemens | MAGNETOM Verio | 3T | 1.0x1.0x1.0 | 3.06 | 1800 | 256 |
| ABIDEI-NYU | Siemens | MAGNETOM Allegra | 3T | 1.3x1.0x1.3 | 3.25 | 2530 | 256 |
| ABIDEI-OHSU | Siemens | MAGNETOM TrioTim | 3T | 1x1x1.1 | 3.58 | 2300 | 256 |
| ABIDEI-OLIN | Siemens | MAGNETOM Allegra | 3T | 1.0x1.0x1.0 | 2.74 | 2500 | 256 |

| Site/Scanner | Vendor | Scanner | Field strength (T) | Voxel Size (mm) | TE (ms) | TR (ms) | FOV (mm) |
|---|---|---|---|---|---|---|---|
| ABIDEI-PITT | Siemens | Allegra | 3T | 1.1x1.1x1.1 | 3.93 | 2100 | 269 |
| ABIDEI-SBL | Philips | Interna | 3T | 1.0x1.0x1.0 | 3.5 | 9 | 256 |
| ABIDEI-SDSU | GE | MR750 | 3T | 1.0x1.0x1.0 | 230 | 2000 | 256 |
| ABIDEI-Stanford | GE | Signa | 3T | .86x1.5x.86 | 1.8 | 8.4 | 220 |
| ABIDEI-Trinity | Philips | Achieva | 3T | 1.0x1.0x1.0 | 3.9 | 8.5 | 256 |
| ABIDEI-UCLA_1 | Siemens | Magnetom TrioTim | 3T | 1.0x1.0x1.2 | 2.84 | 2300 | 256 |
| ABIDEI-UCLA_2 | Siemens | Magnetom TrioTim | 3T | 1.0x1.0x1.2 | 34 | 5000 | 256 |
| ABIDEI-UM_1 | GE | Signa | 3T | 1.2x1.0x1.0 | 250 | 5.7 | 220 |
| ABIDEI-UM_2 | GE | Signa | 3T | 1.0x1.0x1.2 | 250 | 5.7 | 220 |
| ABIDEI-USM | Siemens | Magnetom TrioTim | 3T | 1.0x1.0x1.2 | 2.91 | 2300 | 256 |
| ABIDEI-Yale | Siemens | Magnetom TrioTim | 3T | 1.0x1.0x1.0 | 1.73 | 1230 | 250 |
| ABIDEII-BNI_1 | Philips | Ingenia | 3T | 1.1x1.1x1.2 | 3.1 | 6.7 | 270 |
| ABIDEII-EMC_1 | GE | MR750 | 3T | 0.9x0.9x0.9 | 4.2 | 10.3 | 230 |
| ABIDEII-ETH_1 | Philips | Achieva | 3T | 0.9x0.9x0.9 | 3.9 | 8.4 | 230 |
| ABIDEII-GU_1 | Siemens | TrioTim | 3T | 1.0x1.0x1.0 | 3.5 | 2530 | 256 |
| ABIDEII-IPI_1 | Philips | Achieva | 1.5T | 1.0x1.0x1.0 | 5.6 | 25 | 240 |
| ABIDEII-IU_1 | Siemens | TrioTim | 3T | 0.7x0.7x0.7 | 2.3 | 2400 | 224 |
| ABIDEII-KKI_1 | Philips | Achieva | 3T | 1.0x1.0x1.0 | 3.7 | 8.2/8.0 | 256/212 |
| ABIDEII-KUL_3 | Philips | Achieva Ds | 3T | 1.2x1.2x1.2 | 4.6 | 9.4 | 250 |
| ABIDEII-NYU_1 | Siemens | Allegra | 3T | 1.3x1.0x1.3 | 3.25 | 3.25 | 256 |

| Site/Scanner | Vendor | Scanner | Field strength (T) | Voxel Size (mm) | TE (ms) | TR (ms) | FOV (mm) |
|---|---|---|---|---|---|---|---|
| ABIDEII-OHSU_1 | Siemens | TrioTim | 3T | 1.0x1.0x1.1 | 3.58 | 2300 | 256 |
| ABIDEII-ONRC_2 | Siemens | Skyra | 3T | 0.8x0.8x0.8 | 2.88 | 2200 | 256 |
| ABIDEII-SDSU_1 | GE | MR750 | 3T | 1.0x1.0x1.0 | 3.172 | 8.136 | 256 |
| ABIDEII-STANFORD | GE | Signa | 3T | 0.94x0.94x1.0 | 1.8 | 5.9 | 240 |
| ABIDEII-TCD_1 | Philips | Achieva | 3T | 0.9x0.9x0.9 | 3.9 | 8.4 | 230 |
| ABIDEII-UCD_1 | Siemens | TrioTim | 3T | 1.0x1.0x1.0 | 3.16 | 2000 | 256 |
| ABIDEII-UCLA_1 | Siemens | TrioTim | 3T | 1.0x1.0x1.2 | 2.86 | 2300 | 256 |
| ABIDEII-UM_1 | GE | ** | 3T | 1.0x1.0x1.0 | ** | ** | 256 |
| ABIDEII-USM_1 | Siemens | TrioTim | 3T | 1.0x1.0x1.2 | 2.91 | 900 | 256 |
| CBIC | Siemens | Prisma | 3T | 0.8x0.8x0.8 | 3.15 | 2500 | 224 |
| CUNY | Siemens | Prisma | 3T | 0.8x0.8x0.8 | 3.15 | 2500 | 224 |
| RU | Siemens | TrioTim | 3T | 0.8x0.8x0.8 | 3.15 | 2500 | 224 |
| SI | Siemens | Avanto | 1.5 T | 1.0x1.0x1.0 | 1.64 | 2730 | 176 |

** A double asterisk denotes that a scanning parameter was not able to be identified from existing documentation.

**Table S3 Regions showing lower gray matter volumes in autism**

| Category | ROI | Hemisphere | Voxel Count | Peak T | Peak T MNI coordinates | Peak Std. ß | Peak Std. ß MNI coordinates |
|---|---|---|---|---|---|---|---|
| Cortical | Frontal Medial Cortex | L | 31 | 4.768 | (9, 31, -23) | 0.0739 | (9, 31, -23) |
| | | R | 16 | 3.812 | (-1, 43, -16) | 0.06126 | (-1, 43, -16) |
| | Frontal Operculum Cortex | L | 12 | 4.231 | (34, 25, 7) | 0.06989 | (34, 25, 7) |
| | | R | 12 | 4.318 | (-40, 17, 2) | 0.07169 | (-40, 17, 2) |
| | Frontal Orbital Cortex | L | 559 | 4.919 | (9, 30, -24) | 0.07658 | (11, 12, -22) |
| | | R | 158 | 4.633 | (-20, 27, -22) | 0.07293 | (-20, 27, -22) |
| | Frontal Pole | L | 523 | 5.246 | (29, 44, -16) | 0.08181 | (27, 43, -17) |
| | | R | 546 | 5.171 | (-23, 50, -16) | 0.08283 | (-23, 50, -16) |
| | Inferior Frontal Gyrus, pars opercularis | R | 28 | 4.233 | (-46, 17, 9) | 0.07644 | (-46, 17, 9) |
| | Inferior Frontal Gyrus, pars triangularis | R | 23 | 3.684 | (-54, 32, 13) | 0.06306 | (-39, 30, 19) |
| | Inferior Temporal Gyrus, posterior division | R | 46 | 4.023 | (-49, -43, -19) | 0.07123 | (-52, -43, -21) |
| | Inferior Temporal Gyrus, temporooccipital part | L | 12 | 3.773 | (52, -50, -22) | 0.06183 | (52, -50, -22) |
| | | R | 19 | 4.034 | (-52, -44, -21) | 0.07229 | (-52, -44, -21) |
| | Lateral Occipital Cortex, superior division | L | 197 | 4.795 | (44, -76, 31) | 0.07925 | (44, -76, 31) |
| | | R | 123 | 4.349 | (-46, -73, 36) | 0.07488 | (-47, -73, 34) |
| | Lingual Gyrus | R | 20 | 4.201 | (-19, -49, -13) | 0.0687 | (-19, -49, -13) |
| | Middle Frontal Gyrus | R | 18 | 3.998 | (-42, 10, 30) | 0.06812 | (-42, 10, 30) |
| | Occipital Pole | L | 10 | 3.773 | (39, -90, 10) | 0.06372 | (39, -90, 10) |
| | | R | 14 | 4.158 | (-18, -97, 9) | 0.07573 | (-17, -97, 10) |
| | Paracingulate Gyrus | L | 46 | 4.416 | (10, 39, -7) | 0.0723 | (10, 39, -7) |
| | Parahippocampal Gyrus, anterior division | L | 255 | 5.221 | (23, 2, -22) | 0.07366 | (19, 5, -27) |

| Category | ROI | Hemisphere | Voxel Count | Peak T | Peak T MNI coordinates | Peak Std. ß | Peak Std. ß MNI coordinates |
|---|---|---|---|---|---|---|---|
| | | R | 163 | 5.027 | (-20, -2, -25) | 0.07169 | (-20, -2, -25) |
| | Parahippocampal Gyrus, posterior division | R | 21 | 3.577 | (-20, -37, -16) | 0.05945 | (-20, -37, -17) |
| | Parietal Operculum Cortex | L | 52 | 4.501 | (47, -30, 19) | 0.07385 | (47, -30, 19) |
| | Precentral Gyrus | L | 40 | 3.994 | (2, -28, 72) | 0.06661 | (45, 5, 35) |
| | Subcallosal Cortex | L | 195 | 4.787 | (9, 30, -23) | 0.0736 | (9, 30, -23) |
| | | R | 26 | 3.479 | (-11, 21, -20) | 0.05274 | (-8, 24, -23) |
| | Superior Frontal Gyrus | R | 19 | 3.783 | (-22, 3, 54) | 0.06446 | (-22, 3, 54) |
| | Temporal Fusiform Cortex, anterior division | R | 28 | 3.844 | (-24, 0, -45) | 0.0638 | (-23, 0, -45) |
| | Temporal Pole | L | 144 | 4.629 | (25, 4, -25) | 0.06848 | (21, 5, -40) |
| | | R | 24 | 3.911 | (-24, 3, -24) | 0.05675 | (-23, 3, -24) |
| Subcortical | Accumbens | L | 14 | 3.669 | (-13, 7, -12) | 0.0502 | (-11, 6, -9) |
| | Amygdala | L | 1093 | 5.796 | (-24, -6, -18) | 0.07679 | (-24, -6, -18) |
| | | R | 1185 | 6.3 | (27, 0, -22) | 0.08736 | (27, 0, -22) |
| | Hippocampus | L | 81 | 5.133 | (-23, -9, -19) | 0.06707 | (-23, -9, -19) |
| | | R | 84 | 5.273 | (23, -6, -23) | 0.07136 | (23, -6, -23) |
| | Putamen | L | 64 | 4.022 | (-14, 6, -12) | 0.05466 | (-16, 5, -11) |
| | | R | 77 | 3.873 | (20, 7, -12) | 0.0541 | (20, 7, -12) |
| | Thalamus | L | 470 | 5.978 | (0, -19, 5) | 0.09839 | (0, -13, 0) |
| | | R | 910 | 7.393 | (1, -19, 5) | 0.126 | (1, -19, 5) |
| Cerebellum | Crus I | L | 768 | 4.997 | (-26, -74, -37) | 0.08285 | (-26, -74, -37) |
| | | R | 669 | 5.332 | (29, -74, -36) | 0.08794 | (29, -74, -36) |
| | Crus II | L | 878 | 5.281 | (-26, -75, -39) | 0.08734 | (-26, -75, -39) |

| Category | ROI | Hemisphere | Voxel Count | Peak T | Peak T MNI coordinates | Peak Std. *ß* | Peak Std. *ß* MNI coordinates |
|---|---|---|---|---|---|---|---|
| | | R | 909 | 5.732 | (30, -70, -40) | 0.09381 | (30, -70, -40) |
| | V | L | 12 | 4.057 | (-26, -31, -30) | 0.07004 | (-26, -31, -30) |
| | VI | L | 665 | 4.762 | (-32, -56, -31) | 0.07135 | (-32, -56, -31) |
| | | R | 163 | 4.328 | (29, -67, -27) | 0.06994 | (39, -36, -34) |
| | VIIIa | L | 620 | 4.907 | (-33, -50, -57) | 0.0804 | (-33, -50, -57) |
| | | R | 474 | 4.589 | (23, -57, -55) | 0.07481 | (26, -52, -57) |
| | VIIIb | L | 27 | 3.88 | (-26, -49, -56) | 0.06617 | (-26, -49, -56) |
| | | R | 395 | 4.772 | (26, -48, -55) | 0.0788 | (26, -49, -55) |
| | VIIb | L | 315 | 4.863 | (-35, -40, -46) | 0.07765 | (-39, -53, -52) |
| | | R | 282 | 4.595 | (21, -71, -50) | 0.07416 | (21, -71, -50) |
| | X | L | 21 | 3.596 | (-23, -39, -48) | 0.06119 | (-23, -39, -48) |

**Table S4 Regions showing lower gray matter amygdalar and thalamic subnuclei volume in autism**

| Category | ROI | Hemisphere | Voxel Count | Peak T | Peak T MNI coordinates | Peak Std. ß | Peak Std. ß MNI coordinates |
|---|---|---|---|---|---|---|---|
| Amygdala | Accessory Basal | L | 114 | 5.217 | (21, -3, -19) | 0.06807 | (21, -3, -19) |
| | | R | 118 | 5.546 | (-21, -4, -17) | 0.07364 | (-21, -4, -18) |
| | Anterior amygdaloid area | L | 26 | 5.623 | (26, 0, -18) | 0.07518 | (26, 0, -18) |
| | | R | 28 | 5.106 | (-24, -4, -15) | 0.06754 | (-24, -4, -15) |
| | Basolateral | L | 211 | 6.091 | (24, -1, -22) | 0.08391 | (24, -1, -22) |
| | | R | 212 | 5.796 | (-24, -6, -18) | 0.07679 | (-24, -6, -18) |
| | Intercalated nuclei | L | 58 | 6.05 | (27, -1, -20) | 0.08331 | (27, -1, -20) |
| | | R | 55 | 5.563 | (-23, -4, -17) | 0.07399 | (-25, -5, -17) |
| | Lateral | L | 185 | 6.3 | (27, 0, -22) | 0.08736 | (27, 0, -22) |
| | | R | 162 | 5.511 | (-27, -4, -22) | 0.07518 | (-26, -4, -22) |
| | Medial, cortical, nucleus of the lateral olfactory tract, amygdalohippocampal area | L | 77 | 4.438 | (20, -5, -12) | 0.05816 | (20, -6, -10) |
| | | R | 84 | 5.454 | (-20, -3, -17) | 0.07119 | (-20, -3, -17) |
| | Paralaminar | L | 68 | 5.233 | (22, -6, -22) | 0.0703 | (21, -1, -24) |
| | | R | 76 | 5.195 | (-20, -3, -25) | 0.07412 | (-20, -3, -25) |
| | Periamygdaloid cortex | R | 21 | 4.42 | (-15, -4, -20) | 0.0583 | (-15, -4, -20) |
| Thalamus | Anterior | L | 77 | 5.552 | (0, -6, 2) | 0.09628 | (0, -6, 2) |
| | | R | 125 | 5.652 | (1, -5, 1) | 0.1008 | (1, -5, 1) |
| | Medio Dorsal | L | 136 | 5.061 | (-1, -12, 2) | 0.07923 | (-1, -12, 1) |
| | | R | 654 | 7.393 | (1, -19, 5) | 0.126 | (1, -19, 5) |
| | Ventral Anterior | L | 85 | 3.879 | (-1, -11, 10) | 0.05991 | (-1, -11, 10) |
| | | R | 267 | 4.401 | (2, -10, 11) | 0.07125 | (2, -10, 11) |

**Table S5 Regions showing lower cerebellar gray matter volume in autism based on the functional parcellation**

| Category | ROI | Hemisphere | Voxel Count | Peak T | Peak T MNI coordinates | Peak Std. *ß* | Peak Std. *ß* MNI coordinates |
|---|---|---|---|---|---|---|---|
| Cerebellum functional parcellation | A1 | L | 12 | 3.58 | (-11, -77, -47) | 0.06236 | (-11, -77, -47) |
| | | R | 21 | 3.967 | (16, -72, -47) | 0.06672 | (16, -72, -47) |
| | A2 | L | 427 | 4.907 | (-33, -50, -57) | 0.0804 | (-33, -50, -57) |
| | | R | 69 | 4.385 | (35, -43, -49) | 0.07462 | (35, -43, -49) |
| | A3 | L | 213 | 4.596 | (-28, -49, -55) | 0.07599 | (-27, -49, -55) |
| | | R | 229 | 4.772 | (26, -48, -55) | 0.0788 | (26, -49, -55) |
| | D1 | L | 230 | 4.272 | (-29, -68, -27) | 0.06928 | (-36, -67, -49) |
| | | R | 10 | 3.643 | (38, -65, -44) | 0.06365 | (39, -65, -44) |
| | D2 | L | 97 | 4.749 | (-43, -44, -45) | 0.07903 | (-43, -44, -45) |
| | D3 | L | 485 | 4.762 | (-32, -56, -31) | 0.07765 | (-39, -53, -52) |
| | | R | 409 | 4.595 | (21, -71, -50) | 0.07416 | (21, -71, -50) |
| | D4 | L | 424 | 4.734 | (-33, -57, -31) | 0.07078 | (-33, -57, -31) |
| | | R | 298 | 5.511 | (31, -69, -40) | 0.09094 | (31, -69, -40) |
| | M2 | L | 40 | 4.863 | (-35, -40, -46) | 0.07705 | (-35, -40, -46) |
| | | R | 27 | 3.929 | (38, -36, -34) | 0.06677 | (38, -36, -34) |
| | M3 | L | 83 | 3.881 | (-23, -58, -54) | 0.06218 | (-23, -58, -54) |
| | | R | 205 | 4.668 | (26, -47, -55) | 0.07583 | (26, -47, -55) |
| | M4 | R | 95 | 4.194 | (25, -48, -55) | 0.06842 | (25, -48, -55) |
| | S1 | L | 483 | 4.902 | (-29, -69, -37) | 0.08068 | (-29, -69, -37) |
| | | R | 613 | 5.569 | (29, -70, -40) | 0.09068 | (29, -70, -40) |
| | S2 | L | 535 | 5.281 | (-26, -75, -39) | 0.08734 | (-26, -75, -39) |

| Category | ROI | Hemisphere | Voxel Count | Peak T | Peak T MNI coordinates | Peak Std. ß | Peak Std. ß MNI coordinates |
|---|---|---|---|---|---|---|---|
| | | R | 323 | 5.695 | (29, -71, -40) | 0.09314 | (29, -71, -40) |
| | S3 | L | 216 | 4.755 | (-27, -75, -36) | 0.07803 | (-27, -75, -36) |
| | | R | 476 | 5.732 | (30, -70, -40) | 0.09381 | (30, -70, -40) |
| | S4 | L | 35 | 3.979 | (-48, -55, -49) | 0.06555 | (-48, -55, -49) |
| | | R | 15 | 3.912 | (17, -70, -55) | 0.0646 | (49, -42, -38) |
| | S5 | L | 10 | 3.604 | (-34, -40, -47) | 0.06201 | (-34, -40, -47) |
| | | R | 96 | 4.579 | (26, -45, -56) | 0.07259 | (39, -43, -49) |

**Table S6 Regions showing higher gray matter volumes in autism**

| Category | ROI | Hemisphere | Voxel Count | Peak T | Peak T MNI coordinates | Peak Std. ß | Peak Std. ß MNI coordinates |
|---|---|---|---|---|---|---|---|
| Cortical | Heschl's Gyrus | L | 22 | -4.294 | (52, -12, 6) | -0.072087 | (52, -12, 6) |
| | Planum Polare | L | 125 | -4.582 | (50, -2, -4) | -0.075104 | (50, -2, -4) |
| | | R | 17 | -5.420 | (-59, -3, 0) | -0.092134 | (-59, -3, 0) |
| | Superior Temporal Gyrus, anterior division | L | 72 | -4.322 | (64, 0, 0) | -0.075443 | (60, 3, -5) |
| | | R | 19 | -5.653 | (-59, -4, 0) | -0.099601 | (-59, -4, 0) |
| Subcortical | Thalamus | R | 12 | -5.768 | (17, -27, 15) | -0.099427 | (17, -27, 15) |
| Thalamus | Ventral Latero Dorsal | R | 12 | -5.768 | (17, -27, 15) | -0.099427 | (17, -27, 15) |

**Table S7 Regions showing lower gray matter volumes in autism covarying FSIQ**

| Category | ROI | Hemisphere | Voxel Count | Peak T | Peak T MNI coordinates | Peak Std. ß | Peak Std. ß MNI coordinates |
|---|---|---|---|---|---|---|---|
| Cortical | Frontal Medial Cortex | L | 19 | 4.544 | (9, 32, -24) | 0.07535 | (9, 32, -24) |
| | Frontal Orbital Cortex | L | 69 | 4.647 | (9, 31, -24) | 0.07716 | (9, 31, -24) |
| | Frontal Pole | L | 128 | 4.852 | (28, 44, -16) | 0.08081 | (28, 44, -16) |
| | | R | 163 | 4.553 | (-20, 54, -18) | 0.08026 | (-20, 54, -18) |
| | Lateral Occipital Cortex, superior division | L | 54 | 4.232 | (43, -78, 32) | 0.07583 | (43, -78, 32) |
| | Paracingulate Gyrus | L | 26 | 3.909 | (10, 39, -8) | 0.06959 | (11, 39, -8) |
| | Parahippocampal Gyrus, anterior division | L | 108 | 4.507 | (23, 2, -24) | 0.06937 | (18, 5, -27) |
| | Parietal Operculum Cortex | L | 14 | 4.394 | (46, -30, 18) | 0.07788 | (46, -30, 18) |
| | Subcallosal Cortex | L | 62 | 4.306 | (10, 19, -21) | 0.06832 | (9, 30, -23) |
| Subcortical | Amygdala | L | 602 | 4.923 | (-24, -5, -18) | 0.07009 | (-24, -5, -18) |
| | | R | 607 | 5.593 | (29, -1, -21) | 0.08209 | (29, -1, -21) |
| | Thalamus | L | 94 | 5.202 | (0, -13, 0) | 0.09943 | (0, -13, -1) |
| | | R | 189 | 7.069 | (1, -19, 5) | 0.12877 | (1, -19, 5) |
| Cerebellum | Crus I | L | 226 | 4.663 | (-26, -75, -37) | 0.08283 | (-26, -75, -37) |
| | | R | 228 | 4.947 | (30, -70, -38) | 0.08697 | (30, -70, -38) |
| | Crus II | L | 303 | 4.941 | (-26, -76, -39) | 0.08657 | (-26, -76, -39) |
| | | R | 422 | 5.341 | (30, -70, -40) | 0.09339 | (30, -70, -40) |
| | VI | L | 30 | 4.054 | (-19, -61, -15) | 0.06647 | (-20, -61, -15) |
| | VIIIb | R | 21 | 3.744 | (26, -49, -56) | 0.0674 | (27, -48, -55) |
| | VIIb | L | 12 | 3.72 | (-39, -53, -53) | 0.06643 | (-39, -53, -52) |

**Table S8 Regions showing lower gray matter amygdalar and thalamic subnuclei volume in autism covarying FSIQ**

| Category | ROI | Hemisphere | Voxel Count | Peak T | Peak T MNI coordinates | Peak Std. ß | Peak Std. ß MNI coordinates |
|---|---|---|---|---|---|---|---|
| Amygdala | Accessory Basal | L | 26 | 4.224 | (21, -3, -19) | 0.05939 | (21, -3, -19) |
| | | R | 89 | 4.779 | (-21, -4, -18) | 0.06762 | (-21, -4, -18) |
| | Anterior amygdaloid area | L | 11 | 4.669 | (26, 0, -18) | 0.06661 | (26, 0, -18) |
| | | R | 19 | 4.123 | (-24, -4, -15) | 0.05884 | (-26, -2, -18) |
| | Basolateral | L | 137 | 5.197 | (24, -1, -22) | 0.07623 | (24, -1, -22) |
| | | R | 146 | 4.923 | (-24, -5, -18) | 0.07009 | (-24, -5, -18) |
| | Intercalated nuclei | L | 50 | 5.22 | (27, -1, -20) | 0.07675 | (27, -1, -20) |
| | | R | 44 | 4.736 | (-23, -4, -17) | 0.06732 | (-25, -4, -18) |
| | Lateral | L | 120 | 5.593 | (29, -1, -21) | 0.08209 | (29, -1, -21) |
| | | R | 77 | 4.595 | (-25, -5, -20) | 0.0664 | (-25, -5, -20) |
| | Medial, cortical, nucleus of the lateral olfactory tract, amygdalohippocampal area | R | 48 | 4.564 | (-20, -3, -17) | 0.06321 | (-20, -3, -17) |
| | Paralaminar | L | 12 | 4.168 | (21, -1, -24) | 0.0618 | (21, -1, -24) |
| | | R | 29 | 4.396 | (-19, -3, -24) | 0.06575 | (-19, -3, -24) |
| Thalamus | Anterior | L | 16 | 4.655 | (0, -6, 2) | 0.08707 | (0, -6, 2) |
| | | R | 52 | 5.228 | (1, -5, 1) | 0.09892 | (1, -5, 1) |
| | Medio Dorsal | R | 213 | 7.069 | (1, -19, 5) | 0.12877 | (1, -19, 5) |

**Table S9 Regions showing lower cerebellar gray matter volume in autism based on the functional parcellation covarying FSIQ**

| Category | ROI | Hemisphere | Voxel Count | Peak T | Peak T MNI coordinates | Peak Std. ß | Peak Std. ß MNI coordinates |
|---|---|---|---|---|---|---|---|
| Cerebellum functional parcellation | A2 | L | 12 | 3.906 | (-20, -64, -17) | 0.06333 | (-20, -62, -15) |
| | A3 | R | 11 | 3.744 | (26, -49, -56) | 0.0674 | (27, -48, -55) |
| | D3 | L | 11 | 3.72 | (-39, -53, -53) | 0.06643 | (-39, -53, -52) |
| | D4 | R | 23 | 5.089 | (31, -69, -40) | 0.08907 | (31, -69, -40) |
| | M3 | L | 13 | 4.054 | (-19, -61, -15) | 0.06647 | (-20, -61, -15) |
| | S1 | L | 145 | 4.317 | (-23, -70, -38) | 0.07783 | (-23, -70, -38) |
| | | R | 235 | 5.149 | (29, -70, -40) | 0.0899 | (29, -70, -40) |
| | S2 | L | 335 | 4.941 | (-26, -76, -39) | 0.08657 | (-26, -76, -39) |
| | | R | 166 | 5.239 | (29, -71, -40) | 0.09184 | (29, -71, -40) |
| | S3 | L | 44 | 4.374 | (-27, -76, -36) | 0.07639 | (-27, -76, -36) |
| | | R | 226 | 5.341 | (30, -70, -40) | 0.09339 | (30, -70, -40) |

#### Table S10 Regions showing higher gray matter volumes in autism covarying FSIQ

| Category | ROI | Hemisphere | Voxel Count | Peak T | Peak T MNI coordinates | Peak Std. ß | Peak Std. ß MNI coordinates |
|---|---|---|---|---|---|---|---|
| Cortical | Planum Polare | L | 32 | -4.164 | (45, -4, -14) | -0.068794 | (45, -4, -15) |
| | | R | 16 | -4.882 | (-59, -3, 0) | -0.088699 | (-59, -3, 0) |
| | Superior Temporal Gyrus, anterior division | L | 10 | -3.900 | (64, 1, 0) | -0.067916 | (64, 1, 0) |
| | | R | 10 | -5.252 | (-60, -6, 2) | -0.095736 | (-60, -6, 2) |

**Table S11 Regions showing lower gray matter volumes in autism excluding mild-moderate motion**

| Category | ROI | Hemisphere | Voxel Count | Peak T | Peak T MNI coordinates | Peak Std. ß | Peak Std. ß MNI coordinates |
|---|---|---|---|---|---|---|---|
| Cortical | Frontal Medial Cortex | L | 17 | 4.063 | (9, 31, -23) | 0.06197 | (9, 32, -24) |
| | Frontal Orbital Cortex | L | 147 | 4.341 | (11, 12, -22) | 0.07473 | (25, 9, -21) |
| | | R | 26 | 3.805 | (-22, 30, -22) | 0.05904 | (-23, 31, -22) |
| | Frontal Pole | L | 86 | 4.525 | (25, 44, -17) | 0.07322 | (25, 44, -18) |
| | | R | 171 | 4.739 | (-23, 50, -16) | 0.07824 | (-22, 51, -13) |
| | Lateral Occipital Cortex, superior division | L | 22 | 4.702 | (44, -76, 31) | 0.07611 | (44, -76, 31) |
| | | R | 16 | 3.943 | (-25, -84, 18) | 0.07002 | (-26, -83, 18) |
| | Paracingulate Gyrus | L | 13 | 3.91 | (10, 39, -7) | 0.06276 | (10, 39, -8) |
| | Parahippocampal Gyrus, anterior division | L | 122 | 4.465 | (23, 2, -22) | 0.07204 | (18, 5, -27) |
| | Parietal Operculum Cortex | L | 23 | 4.54 | (47, -30, 19) | 0.07439 | (47, -30, 19) |
| | Precentral Gyrus | L | 10 | 4.006 | (45, 5, 35) | 0.06885 | (45, 5, 35) |
| | Subcallosal Cortex | L | 80 | 4.212 | (10, 18, -20) | 0.0608 | (10, 18, -21) |
| Subcortical | Amygdala | L | 676 | 4.975 | (-25, -6, -18) | 0.06581 | (-25, -5, -18) |
| | | R | 672 | 5.436 | (28, 0, -22) | 0.0758 | (28, 0, -22) |
| | Thalamus | L | 271 | 5.966 | (0, -19, 5) | 0.10005 | (0, -19, 5) |
| | | R | 445 | 7.206 | (1, -19, 5) | 0.12376 | (1, -19, 5) |
| Cerebellum | Crus I | L | 333 | 4.408 | (-26, -74, -37) | 0.07724 | (-29, -69, -37) |
| | | R | 293 | 4.737 | (30, -70, -38) | 0.08026 | (30, -70, -38) |
| | Crus II | L | 373 | 4.799 | (-26, -75, -39) | 0.08002 | (-26, -75, -39) |
| | | R | 489 | 5.365 | (29, -71, -40) | 0.08894 | (29, -71, -40) |

| Category | ROI | Hemisphere | Voxel Count | Peak T | Peak T MNI coordinates | Peak Std. *ß* | Peak Std. *ß* MNI coordinates |
|---|---|---|---|---|---|---|---|
| | VI | L | 184 | 4.177 | (-32, -56, -31) | 0.06498 | (-36, -52, -31) |
| | | R | 18 | 3.632 | (28, -67, -28) | 0.05575 | (28, -67, -28) |
| | VIIIa | L | 183 | 4.778 | (-33, -50, -57) | 0.0765 | (-33, -50, -57) |
| | | R | 74 | 4.296 | (23, -57, -55) | 0.07186 | (26, -52, -57) |
| | VIIIb | R | 164 | 4.567 | (26, -48, -55) | 0.07658 | (26, -48, -55) |
| | VIIb | L | 45 | 4.108 | (-38, -53, -51) | 0.06886 | (-39, -53, -52) |
| | | R | 84 | 4.435 | (21, -71, -50) | 0.0712 | (39, -43, -49) |

**Table S12 Regions showing lower gray matter amygdalar and thalamic subnuclei volume in autism excluding mild-moderate motion**

| Category | ROI | Hemisphere | Voxel Count | Peak T | Peak T MNI coordinates | Peak Std. ß | Peak Std. ß MNI coordinates |
|---|---|---|---|---|---|---|---|
| Amygdala | Accessory Basal | L | 32 | 4.469 | (21, -3, -19) | 0.05822 | (21, -3, -19) |
| | | R | 84 | 4.521 | (-21, -4, -18) | 0.05969 | (-21, -4, -19) |
| | Anterior amygdaloid area | L | 13 | 4.706 | (26, 0, -18) | 0.06306 | (26, 0, -18) |
| | | R | 23 | 4.367 | (-24, -4, -15) | 0.05892 | (-25, -4, -15) |
| | Basolateral | L | 151 | 5.231 | (23, -1, -22) | 0.07148 | (23, -1, -22) |
| | | R | 179 | 4.975 | (-25, -6, -18) | 0.06581 | (-25, -5, -18) |
| | Intercalated nuclei | L | 52 | 5.243 | (27, -1, -20) | 0.07208 | (27, -1, -20) |
| | | R | 48 | 4.888 | (-25, -5, -17) | 0.06536 | (-25, -5, -17) |
| | Lateral | L | 114 | 5.436 | (28, 0, -22) | 0.0758 | (28, 0, -22) |
| | | R | 84 | 4.477 | (-27, -4, -21) | 0.05978 | (-26, -4, -21) |
| | Medial, cortical, nucleus of the lateral olfactory tract, amygdalohippocampal area | L | 17 | 3.922 | (20, -5, -12) | 0.05653 | (20, -6, -10) |
| | | R | 43 | 4.371 | (-20, -3, -17) | 0.05672 | (-16, -2, -19) |
| | Paralaminar | L | 14 | 4.559 | (22, -6, -22) | 0.062 | (22, -6, -22) |
| | | R | 34 | 4.3 | (-19, -3, -23) | 0.06058 | (-20, -3, -25) |
| Thalamus | Anterior | L | 64 | 5.388 | (0, -6, 2) | 0.09515 | (0, -6, 2) |
| | | R | 110 | 5.531 | (1, -5, 1) | 0.09857 | (1, -5, 1) |
| | Medio Dorsal | L | 47 | 4.797 | (-1, -12, 1) | 0.07752 | (-1, -12, 1) |
| | | R | 459 | 7.206 | (1, -19, 5) | 0.12376 | (1, -19, 5) |
| | Ventral Anterior | L | 12 | 3.734 | (-10, -11, 14) | 0.05441 | (-10, -11, 14) |

**Table S13 Regions showing lower cerebellar gray matter volume in autism based on the functional parcellation excluding mild-moderate motion**

| Category | ROI | Hemisphere | Voxel Count | Peak T | Peak T MNI coordinates | Peak Std. *ß* | Peak Std. *ß* MNI coordinates |
|---|---|---|---|---|---|---|---|
| Cerebellum functional parcellation | A2 | L | 156 | 4.778 | (-33, -50, -57) | 0.0765 | (-33, -50, -57) |
| | | R | 22 | 4.066 | (35, -43, -49) | 0.06969 | (35, -43, -49) |
| | A3 | L | 38 | 4.42 | (-28, -49, -55) | 0.07439 | (-28, -49, -55) |
| | | R | 102 | 4.567 | (26, -48, -55) | 0.07658 | (26, -48, -55) |
| | D1 | L | 28 | 3.833 | (-28, -65, -27) | 0.05996 | (-30, -67, -26) |
| | D2 | L | 33 | 4.661 | (-42, -44, -45) | 0.07901 | (-41, -43, -43) |
| | D3 | L | 125 | 4.177 | (-32, -56, -31) | 0.06886 | (-39, -53, -52) |
| | | R | 70 | 4.435 | (21, -71, -50) | 0.07078 | (21, -71, -50) |
| | D4 | L | 96 | 4.061 | (-33, -57, -31) | 0.06259 | (-33, -57, -31) |
| | | R | 47 | 5.045 | (31, -69, -40) | 0.08371 | (31, -69, -40) |
| | M3 | R | 63 | 4.526 | (26, -47, -55) | 0.07481 | (26, -47, -55) |
| | M4 | R | 27 | 4.068 | (25, -48, -55) | 0.06737 | (25, -48, -55) |
| | S1 | L | 211 | 4.373 | (-29, -69, -37) | 0.07724 | (-29, -69, -37) |
| | | R | 336 | 5.27 | (29, -70, -40) | 0.08683 | (29, -70, -40) |
| | S2 | L | 363 | 4.799 | (-26, -75, -39) | 0.08002 | (-26, -75, -39) |
| | | R | 173 | 5.365 | (29, -71, -40) | 0.08894 | (29, -71, -40) |
| | S3 | L | 68 | 4.209 | (-27, -75, -36) | 0.06973 | (-28, -73, -36) |
| | | R | 248 | 5.327 | (30, -70, -40) | 0.08841 | (30, -70, -40) |
| | S5 | R | 27 | 4.195 | (39, -43, -49) | 0.0712 | (39, -43, -49) |

**Table S14 Regions showing higher gray matter volumes in autism excluding mild-moderate motion**

| Category | ROI | Hemisphere | Voxel Count | Peak T | Peak T MNI coordinates | Peak Std. ß | Peak Std. ß MNI coordinates |
|---|---|---|---|---|---|---|---|
| Cortical | Heschl's Gyrus | L | 11 | -4.215 | (49, -10, 2) | -0.069046 | (49, -10, 2) |
| | Planum Polare | L | 63 | -4.623 | (45, -4, -13) | -0.069343 | (51, -3, -3) |
| | | R | 13 | -5.455 | (-59, -3, 0) | -0.093525 | (-59, -3, 0) |
| | Superior Temporal Gyrus, anterior division | L | 61 | -4.643 | (64, 0, 0) | -0.077344 | (64, 0, 0) |
| | | R | 20 | -5.619 | (-60, -4, 0) | -0.099058 | (-59, -4, 0) |
| Subcortical | Thalamus | R | 10 | -5.158 | (17, -27, 15) | -0.088735 | (17, -27, 15) |
| Thalamus | Ventral Latero Dorsal | R | 10 | -5.158 | (17, -27, 15) | -0.088735 | (17, -27, 15) |

**Table S15 Regions showing lower gray matter volumes in autism excluding any visually detectable motion**

| Category | ROI | Hemisphere | Voxel Count | Peak T | Peak T MNI coordinates | Peak Std. ß | Peak Std. ß MNI coordinates |
|---|---|---|---|---|---|---|---|
| Cortical | Frontal Orbital Cortex | L | 49 | 4.151 | (11, 13, -21) | 0.06721 | (11, 12, -22) |
| | Frontal Pole | R | 24 | 3.99 | (-22, 51, -15) | 0.06878 | (-22, 51, -14) |
| | Parietal Operculum Cortex | L | 14 | 4.532 | (47, -30, 19) | 0.07793 | (47, -30, 19) |
| | Subcallosal Cortex | L | 46 | 4.115 | (10, 19, -21) | 0.0634 | (9, 23, -24) |
| Subcortical | Amygdala | L | 270 | 4.286 | (-25, -6, -18) | 0.06132 | (-16, -2, -18) |
| | | R | 293 | 4.879 | (29, -1, -21) | 0.06946 | (29, -1, -20) |
| | Thalamus | L | 181 | 5.01 | (0, -14, 1) | 0.09125 | (0, -13, 0) |
| | | R | 281 | 6.707 | (1, -19, 5) | 0.11807 | (1, -19, 5) |
| Cerebellum | Crus I | L | 93 | 4.055 | (-28, -74, -38) | 0.06978 | (-29, -68, -38) |
| | | R | 91 | 4.365 | (29, -75, -36) | 0.07475 | (32, -70, -39) |
| | Crus II | L | 118 | 4.303 | (-26, -76, -39) | 0.0758 | (-41, -43, -44) |
| | | R | 168 | 4.537 | (32, -69, -40) | 0.07896 | (32, -69, -40) |
| | VI | L | 55 | 3.917 | (-32, -57, -32) | 0.06232 | (-32, -56, -31) |
| | VIIIa | L | 106 | 4.775 | (-33, -50, -57) | 0.08189 | (-32, -49, -57) |
| | | R | 22 | 4.355 | (26, -52, -57) | 0.07344 | (26, -52, -57) |
| | VIIIb | R | 102 | 4.571 | (26, -51, -57) | 0.07955 | (26, -48, -55) |
| | VIIb | L | 19 | 3.929 | (-39, -53, -52) | 0.07086 | (-39, -53, -52) |
| | | R | 17 | 4.166 | (21, -72, -51) | 0.06927 | (21, -71, -50) |

**Table S16 Regions showing lower gray matter amygdalar and thalamic subnuclei volume in autism excluding any visually detectable motion**

| Category | ROI | Hemisphere | Voxel Count | Peak T | Peak T MNI coordinates | Peak Std. ß | Peak Std. ß MNI coordinates |
|---|---|---|---|---|---|---|---|
| Amygdala | Accessory Basal | R | 29 | 3.982 | (-20, -4, -19) | 0.05349 | (-20, -3, -20) |
| | Basolateral | L | 81 | 4.468 | (24, -1, -22) | 0.06193 | (24, 0, -22) |
| | | R | 90 | 4.286 | (-25, -6, -18) | 0.05757 | (-25, -5, -18) |
| | Intercalated nuclei | L | 34 | 4.558 | (28, -1, -19) | 0.06555 | (28, -1, -19) |
| | | R | 26 | 4.16 | (-25, -5, -17) | 0.05729 | (-25, -4, -18) |
| | Lateral | L | 53 | 4.879 | (29, -1, -21) | 0.06946 | (29, -1, -20) |
| | | R | 27 | 4.099 | (-27, -3, -22) | 0.05694 | (-27, -3, -22) |
| | Paralaminar | R | 11 | 3.865 | (-18, -4, -22) | 0.05263 | (-19, -3, -24) |
| Thalamus | Anterior | L | 46 | 4.773 | (0, -6, 2) | 0.0875 | (0, -6, 2) |
| | | R | 87 | 5.322 | (1, -5, 1) | 0.098 | (1, -5, 1) |
| | Medio Dorsal | L | 19 | 4.391 | (-1, -12, 2) | 0.07272 | (-1, -11, 1) |
| | | R | 307 | 6.707 | (1, -19, 5) | 0.11807 | (1, -19, 5) |

**Table S17 Regions showing lower cerebellar gray matter volume in autism based on the functional parcellation excluding any visually detectable motion**

| Category | ROI | Hemisphere | Voxel Count | Peak T | Peak T MNI coordinates | Peak Std. *ß* | Peak Std. *ß* MNI coordinates |
|---|---|---|---|---|---|---|---|
| Cerebellum functional parcellation | A2 | L | 88 | 4.775 | (-33, -50, -57) | 0.08189 | (-32, -49, -57) |
| | A3 | L | 21 | 4.488 | (-28, -49, -55) | 0.07428 | (-28, -49, -55) |
| | | R | 63 | 4.571 | (26, -51, -57) | 0.07955 | (26, -48, -55) |
| | D2 | L | 13 | 4.183 | (-42, -43, -44) | 0.0758 | (-41, -43, -44) |
| | D3 | L | 35 | 3.929 | (-39, -53, -52) | 0.07086 | (-39, -53, -52) |
| | | R | 17 | 4.166 | (21, -72, -51) | 0.06927 | (21, -71, -50) |
| | D4 | L | 37 | 3.917 | (-32, -57, -32) | 0.06029 | (-32, -57, -32) |
| | M3 | R | 38 | 4.41 | (26, -47, -55) | 0.07748 | (26, -47, -55) |
| | M4 | R | 12 | 3.793 | (25, -49, -56) | 0.06786 | (19, -56, -58) |
| | S1 | L | 34 | 3.952 | (-29, -69, -38) | 0.06978 | (-29, -68, -38) |
| | | R | 90 | 4.38 | (29, -70, -39) | 0.07561 | (29, -70, -39) |
| | S2 | L | 150 | 4.303 | (-26, -76, -39) | 0.07535 | (-26, -76, -39) |
| | | R | 65 | 4.407 | (29, -71, -39) | 0.0762 | (29, -71, -39) |
| | S3 | L | 13 | 3.893 | (-28, -73, -36) | 0.06591 | (-27, -75, -36) |
| | | R | 100 | 4.537 | (32, -69, -40) | 0.07896 | (32, -69, -40) |

**Table S18 Regions showing higher gray matter volumes in autism excluding any visually detectable motion**

| Category | ROI | Hemisphere | Voxel Count | Peak T | Peak T MNI coordinates | Peak Std. *ß* | Peak Std. *ß* MNI coordinates |
|---|---|---|---|---|---|---|---|
| Cortical | Planum Polare | L | 22 | -4.220 | (45, -5, -13) | -0.071576 | (62, -1, 2) |
| | Superior Temporal Gyrus, anterior division | L | 64 | -5.151 | (64, 1, -1) | -0.085743 | (64, 0, 0) |
| | | R | 13 | -5.418 | (-60, -4, 0) | -0.102071 | (-59, -4, 0) |

**Table S19 Regions showing lower white matter volume in autism**

| Category | ROI | Hemisphere | Voxel Count | Peak T | Peak T MNI coordinates | Peak Std. ß | Peak Std. ß MNI coordinates |
|---|---|---|---|---|---|---|---|
| cerebrum | Anterior corona radiata | L | 6334 | 6.738 | (-20, 38, -7) | 0.07825 | (-21, 37, -7) |
| | | R | 6332 | 6.514 | (20, 37, -7) | 0.07386 | (18, 39, -8) |
| | Anterior limb of internal capsule | L | 263 | 4.645 | (-24, 5, 18) | 0.05465 | (-21, 12, 14) |
| | | R | 88 | 4.826 | (25, 10, 18) | 0.0524 | (22, 16, 16) |
| | Body of corpus callosum | - | 2780 | 5.4 | (20, 16, 26) | 0.06952 | (5, -30, 17) |
| | Cingulum (cingulate gyrus) | L | 98 | 3.209 | (-9, 34, 11) | 0.04147 | (-9, 17, 27) |
| | | R | 244 | 3.787 | (10, 0, 35) | 0.05289 | (10, 27, 21) |
| | External capsule | L | 486 | 4.973 | (-31, -3, 18) | 0.05527 | (-31, -3, 18) |
| | | R | 385 | 4.902 | (27, 10, 18) | 0.05882 | (26, 17, 8) |
| | Fornix (cres) / Stria terminalis | L | 226 | 3.991 | (-25, -31, 3) | 0.056 | (-25, -31, -4) |
| | | R | 173 | 4.566 | (26, -28, -5) | 0.06653 | (26, -28, -5) |
| | Genu of corpus callosum | - | 4604 | 5.61 | (14, 36, -3) | 0.06673 | (14, 36, -3) |
| | Posterior corona radiata | L | 2570 | 4.249 | (-28, -49, 26) | 0.04516 | (-29, -48, 25) |
| | | R | 2626 | 4.105 | (23, -29, 38) | 0.0436 | (20, -28, 37) |
| | Posterior limb of internal capsule | L | 224 | 4.25 | (-27, -10, 18) | 0.04213 | (-26, -9, 18) |
| | | R | 425 | 3.835 | (23, -22, 11) | 0.04995 | (23, -22, 11) |
| | Posterior thalamic radiation | L | 1200 | 4.823 | (-31, -70, 9) | 0.06016 | (-31, -70, 9) |
| | | R | 1797 | 4.621 | (32, -71, 7) | 0.05919 | (30, -58, 12) |
| | Retrolenticular part of internal capsule | L | 1429 | 4.302 | (-34, -34, 2) | 0.04819 | (-35, -35, -2) |
| | | R | 1308 | 4.316 | (33, -31, 2) | 0.04852 | (26, -22, -3) |

| Category | ROI | Hemisphere | Voxel Count | Peak T | Peak T MNI coordinates | Peak Std. ß | Peak Std. ß MNI coordinates |
|---|---|---|---|---|---|---|---|
| | Sagittal stratum | L | 317 | 3.47 | (-37, -33, -4) | 0.0447 | (-38, -37, -7) |
| | | R | 1110 | 4.981 | (38, -40, -6) | 0.0714 | (38, -39, -6) |
| | Splenium of corpus callosum | - | 5622 | 5.077 | (9, -33, 18) | 0.0721 | (9, -33, 18) |
| | Superior corona radiata | L | 6988 | 5.575 | (-29, -2, 21) | 0.0564 | (-24, 15, 30) |
| | | R | 6954 | 5.463 | (27, -5, 23) | 0.06655 | (16, 9, 39) |
| | Superior fronto-occipital fasciculus | L | 238 | 4.769 | (-22, 11, 23) | 0.04901 | (-22, 3, 21) |
| | | R | 193 | 4.91 | (23, -5, 23) | 0.05231 | (22, -5, 23) |
| | Superior longitudinal fasciculus | L | 5149 | 5.506 | (-31, -6, 23) | 0.06292 | (-32, -2, 18) |
| | | R | 5185 | 5.302 | (32, -5, 23) | 0.05301 | (36, -15, 24) |
| | Tapetum | L | 30 | 3.083 | (-26, -51, 20) | 0.03902 | (-24, -45, 21) |
| | | R | 15 | 2.844 | (30, -51, 17) | 0.03357 | (30, -51, 17) |
| | Uncinate fasciculus | R | 10 | 2.573 | (35, 3, -13) | 0.03797 | (35, 3, -14) |
| cerebellum/ brainstem | Cerebral peduncle | L | 172 | 3.92 | (-7, -22, -21) | 0.04614 | (-7, -22, -21) |
| | | R | 250 | 4.209 | (6, -21, -21) | 0.0464 | (6, -20, -21) |
| | Corticospinal tract | L | 1320 | 5.406 | (-6, -24, -39) | 0.06726 | (-7, -24, -38) |
| | | R | 1331 | 5.227 | (7, -23, -38) | 0.06526 | (7, -23, -38) |
| | Inferior cerebellar peduncle | L | 395 | 5.347 | (-8, -40, -40) | 0.07078 | (-8, -40, -40) |
| | | R | 416 | 4.754 | (7, -40, -50) | 0.06276 | (10, -41, -37) |
| | Medial lemniscus | L | 576 | 5.408 | (-6, -35, -39) | 0.06676 | (-6, -35, -39) |
| | | R | 559 | 4.994 | (8, -35, -37) | 0.06193 | (8, -35, -37) |
| | Middle cerebellar peduncle | - | 11553 | 5.96 | (8, -17, -37) | 0.0791 | (8, -17, -37) |

| Category | ROI | Hemisphere | Voxel Count | Peak T | Peak T MNI coordinates | Peak Std. *ß* | Peak Std. *ß* MNI coordinates |
|---|---|---|---|---|---|---|---|
| | Pontine crossing tract (a part of MCP) | - | 1482 | 5.259 | (-7, -33, -37) | 0.06602 | (-7, -33, -37) |
| | Superior cerebellar peduncle | L | 188 | 4.145 | (-6, -31, -21) | 0.04869 | (-6, -31, -21) |
| | | R | 228 | 4.277 | (5, -34, -22) | 0.05119 | (8, -37, -26) |

**Table S20 Regions showing lower white matter volume in autism covarying FSIQ**

| Category | ROI | Hemisphere | Voxel Count | Peak T | Peak T MNI coordinates | Peak Std. ß | Peak Std. ß MNI coordinates |
|---|---|---|---|---|---|---|---|
| cerebrum | Anterior corona radiata | L | 6368 | 7.194 | (-20, 38, -7) | 0.08779 | (-21, 37, -7) |
| | | R | 6325 | 6.62 | (19, 37, -6) | 0.08057 | (14, 32, -12) |
| | Anterior limb of internal capsule | L | 310 | 4.883 | (-24, 4, 18) | 0.05891 | (-22, 12, 14) |
| | | R | 148 | 4.894 | (25, 10, 18) | 0.06 | (22, 16, 16) |
| | Body of corpus callosum | - | 2794 | 5.419 | (-19, 19, 25) | 0.07286 | (15, 14, 23) |
| | Cingulum (cingulate gyrus) | L | 155 | 3.487 | (-11, -50, 17) | 0.05274 | (-7, -43, 29) |
| | | R | 352 | 4.11 | (10, 0, 35) | 0.06192 | (10, 27, 21) |
| | External capsule | L | 421 | 4.917 | (-26, 9, 18) | 0.0565 | (-27, 10, 18) |
| | | R | 387 | 4.924 | (26, 10, 18) | 0.05541 | (26, 17, 8) |
| | Fornix (cres) / Stria terminalis | L | 187 | 3.859 | (-24, -31, 3) | 0.05208 | (-25, -31, -4) |
| | | R | 160 | 4.48 | (25, -30, 2) | 0.06149 | (25, -30, 2) |
| | Genu of corpus callosum | - | 4824 | 6.146 | (14, 36, -3) | 0.07811 | (14, 36, -3) |
| | Posterior corona radiata | L | 2741 | 4.518 | (-28, -49, 26) | 0.0514 | (-28, -49, 26) |
| | | R | 2848 | 4.437 | (32, -56, 19) | 0.0527 | (32, -56, 19) |
| | Posterior limb of internal capsule | L | 187 | 3.845 | (-27, -10, 18) | 0.04004 | (-26, -15, 15) |
| | | R | 534 | 3.955 | (23, -15, 18) | 0.05014 | (23, -22, 11) |
| | Posterior thalamic radiation | L | 1552 | 5.238 | (-31, -70, 9) | 0.06972 | (-31, -70, 9) |
| | | R | 2189 | 5.3 | (32, -70, 9) | 0.06455 | (30, -58, 12) |
| | Retrolenticular part of internal capsule | L | 1148 | 4.058 | (-33, -35, 3) | 0.04746 | (-33, -36, 3) |
| | | R | 1406 | 4.273 | (33, -31, 2) | 0.05041 | (33, -31, 2) |

| Category | ROI | Hemisphere | Voxel Count | Peak T | Peak T MNI coordinates | Peak Std. ß | Peak Std. ß MNI coordinates |
|---|---|---|---|---|---|---|---|
| | Sagittal stratum | L | 548 | 3.632 | (-37, -42, -4) | 0.05632 | (-38, -37, -8) |
| | | R | 1091 | 4.648 | (38, -40, -5) | 0.07145 | (38, -39, -6) |
| | Splenium of corpus callosum | - | 6376 | 4.931 | (11, -43, 20) | 0.07361 | (11, -43, 20) |
| | Superior corona radiata | L | 6898 | 5.864 | (-24, 15, 30) | 0.06553 | (-24, 15, 30) |
| | | R | 7015 | 5.692 | (22, 11, 35) | 0.07247 | (16, 9, 39) |
| | Superior fronto-occipital fasciculus | L | 248 | 4.931 | (-22, 10, 23) | 0.05527 | (-22, 3, 21) |
| | | R | 189 | 5.029 | (23, -5, 23) | 0.05693 | (22, -5, 23) |
| | Superior longitudinal fasciculus | L | 5496 | 5.053 | (-32, -1, 18) | 0.06683 | (-32, -1, 18) |
| | | R | 5558 | 5.039 | (32, -5, 23) | 0.05704 | (44, -25, 29) |
| | Tapetum | L | 32 | 3.702 | (-26, -51, 18) | 0.0491 | (-26, -51, 18) |
| | | R | 31 | 3.432 | (30, -51, 17) | 0.04337 | (30, -51, 17) |
| cerebellum/ brainstem | Cerebral peduncle | L | 38 | 3.223 | (-6, -23, -21) | 0.0404 | (-7, -22, -21) |
| | | R | 179 | 3.384 | (6, -21, -21) | 0.03971 | (6, -21, -21) |
| | Corticospinal tract | L | 1228 | 4.673 | (-7, -19, -29) | 0.0624 | (-7, -28, -39) |
| | | R | 1292 | 4.563 | (4, -28, -37) | 0.05935 | (4, -28, -37) |
| | Inferior cerebellar peduncle | L | 337 | 4.264 | (-9, -39, -41) | 0.05939 | (-8, -40, -40) |
| | | R | 348 | 4.038 | (10, -40, -38) | 0.05617 | (10, -40, -37) |
| | Medial lemniscus | L | 538 | 4.427 | (-6, -35, -39) | 0.05781 | (-6, -35, -39) |
| | | R | 542 | 4.367 | (5, -36, -42) | 0.06173 | (8, -37, -28) |
| | Middle cerebellar peduncle | - | 10837 | 5.273 | (7, -20, -40) | 0.07671 | (29, -50, -42) |
| | Pontine crossing tract (a part of MCP) | - | 1472 | 4.54 | (-7, -33, -37) | 0.06045 | (-7, -33, -37) |

| Category | ROI | Hemisphere | Voxel Count | Peak T | Peak T MNI coordinates | Peak Std. ß | Peak Std. ß MNI coordinates |
|---|---|---|---|---|---|---|---|
| | Superior cerebellar peduncle | L | 98 | 3.457 | (-6, -29, -17) | 0.04314 | (-6, -29, -17) |
| | | R | 164 | 3.621 | (7, -29, -18) | 0.05328 | (8, -37, -26) |

#### Table S21 Regions showing lower white matter volume in autism excluding mild-moderate motion

| Category | ROI | Hemisphere | Voxel Count | Peak T | Peak T MNI coordinates | Peak Std. ß | Peak Std. ß MNI coordinates |
|---|---|---|---|---|---|---|---|
| cerebrum | Anterior corona radiata | L | 6330 | 6.608 | (-20, 38, -7) | 0.07684 | (-21, 37, -7) |
| | | R | 6340 | 6.23 | (20, 38, -7) | 0.07118 | (20, 39, -7) |
| | Anterior limb of internal capsule | L | 231 | 4.593 | (-24, 4, 18) | 0.05385 | (-21, 12, 14) |
| | | R | 92 | 4.638 | (25, 10, 18) | 0.05213 | (22, 16, 16) |
| | Body of corpus callosum | - | 2926 | 5.206 | (20, 18, 25) | 0.06797 | (4, -30, 17) |
| | Cingulum (cingulate gyrus) | L | 132 | 3.455 | (-11, -50, 17) | 0.0462 | (-10, -52, 17) |
| | | R | 254 | 3.657 | (10, 0, 35) | 0.05148 | (10, 28, 20) |
| | External capsule | L | 530 | 5.011 | (-31, -3, 18) | 0.05706 | (-31, -3, 18) |
| | | R | 481 | 4.728 | (29, -5, 18) | 0.05863 | (26, 17, 8) |
| | Fornix (cres) / Stria terminalis | L | 247 | 4.146 | (-24, -31, -3) | 0.05856 | (-25, -31, -4) |
| | | R | 182 | 4.597 | (26, -28, -5) | 0.06735 | (26, -28, -5) |
| | Genu of corpus callosum | - | 4581 | 5.497 | (14, 36, -3) | 0.06571 | (14, 36, -3) |
| | Posterior corona radiata | L | 2640 | 4.525 | (-29, -49, 25) | 0.04846 | (-29, -49, 25) |
| | | R | 2712 | 4.108 | (27, -45, 30) | 0.04461 | (33, -57, 19) |
| | Posterior limb of internal capsule | L | 311 | 4.29 | (-27, -10, 18) | 0.04333 | (-26, -9, 18) |
| | | R | 608 | 4.096 | (23, -22, 11) | 0.05341 | (23, -22, 11) |
| | Posterior thalamic radiation | L | 1262 | 4.729 | (-31, -70, 9) | 0.05952 | (-31, -70, 9) |
| | | R | 1786 | 4.79 | (32, -71, 7) | 0.06128 | (30, -58, 12) |
| | Retrolenticular part of internal capsule | L | 1600 | 4.331 | (-33, -34, 3) | 0.04971 | (-35, -35, -2) |
| | | R | 1479 | 4.418 | (33, -31, 2) | 0.05165 | (26, -22, -3) |

| Category | ROI | Hemisphere | Voxel Count | Peak T | Peak T MNI coordinates | Peak Std. ß | Peak Std. ß MNI coordinates |
|---|---|---|---|---|---|---|---|
| | Sagittal stratum | L | 314 | 3.501 | (-37, -32, -4) | 0.04628 | (-38, -37, -7) |
| | | R | 1129 | 4.905 | (38, -39, -5) | 0.07213 | (38, -39, -6) |
| | Splenium of corpus callosum | - | 6032 | 4.944 | (11, -42, 21) | 0.07328 | (-4, -38, 22) |
| | Superior corona radiata | L | 7027 | 5.454 | (-28, -6, 23) | 0.05637 | (-24, 15, 30) |
| | | R | 6987 | 5.366 | (21, 10, 36) | 0.06737 | (16, 9, 39) |
| | Superior fronto-occipital fasciculus | L | 239 | 4.702 | (-22, 11, 23) | 0.04863 | (-22, 11, 23) |
| | | R | 196 | 4.775 | (23, -5, 23) | 0.0515 | (22, -5, 23) |
| | Superior longitudinal fasciculus | L | 5210 | 5.421 | (-31, -2, 20) | 0.06728 | (-32, -2, 18) |
| | | R | 5249 | 5.267 | (34, -6, 23) | 0.0528 | (36, -15, 24) |
| | Tapetum | L | 30 | 3.387 | (-26, -51, 20) | 0.04265 | (-24, -45, 21) |
| | | R | 25 | 3.024 | (30, -51, 17) | 0.03612 | (30, -51, 17) |
| cerebellum/ brainstem | Cerebral peduncle | L | 98 | 3.408 | (-7, -23, -21) | 0.04008 | (-7, -22, -21) |
| | | R | 138 | 3.378 | (6, -21, -21) | 0.03745 | (8, -26, -19) |
| | Corticospinal tract | L | 1235 | 4.343 | (-7, -19, -28) | 0.05466 | (-7, -28, -39) |
| | | R | 1252 | 4.149 | (7, -24, -39) | 0.05197 | (7, -25, -40) |
| | Inferior cerebellar peduncle | L | 348 | 4.75 | (-8, -40, -40) | 0.06236 | (-8, -40, -40) |
| | | R | 377 | 4.003 | (8, -41, -43) | 0.05174 | (10, -41, -37) |
| | Medial lemniscus | L | 541 | 4.396 | (-6, -35, -39) | 0.05705 | (-7, -39, -40) |
| | | R | 541 | 4.334 | (8, -37, -28) | 0.05779 | (8, -37, -28) |
| | Middle cerebellar peduncle | - | 10912 | 5.232 | (29, -51, -43) | 0.07275 | (29, -51, -43) |
| | Pontine crossing tract (a part of MCP) | - | 1466 | 4.147 | (-6, -33, -38) | 0.05193 | (-7, -33, -37) |

| Category | ROI | Hemisphere | Voxel Count | Peak T | Peak T MNI coordinates | Peak Std. *ß* | Peak Std. *ß* MNI coordinates |
|---|---|---|---|---|---|---|---|
| | Superior cerebellar peduncle | L | 144 | 3.516 | (-6, -31, -21) | 0.0416 | (-6, -31, -21) |
| | | R | 175 | 3.798 | (5, -34, -22) | 0.04819 | (8, -37, -26) |

**Table S22 Regions showing lower white matter volume in autism excluding any visually detectable motion**

| Category | ROI | Hemisphere | Voxel Count | Peak T | Peak T MNI coordinates | Peak Std. ß | Peak Std. ß MNI coordinates |
|---|---|---|---|---|---|---|---|
| cerebrum | Anterior corona radiata | L | 6169 | 5.781 | (-19, 39, -7) | 0.06933 | (-17, 39, -8) |
| | | R | 6265 | 5.673 | (20, 39, -7) | 0.06753 | (20, 39, -7) |
| | Anterior limb of internal capsule | L | 174 | 3.987 | (-24, 4, 18) | 0.05015 | (-21, 11, 14) |
| | | R | 71 | 3.944 | (25, 10, 17) | 0.04711 | (22, 16, 16) |
| | Body of corpus callosum | | 2150 | 4.473 | (20, 16, 26) | 0.06276 | (4, -30, 17) |
| | Cingulum (cingulate gyrus) | L | 85 | 3.116 | (-7, -30, 32) | 0.04418 | (-10, -52, 18) |
| | | R | 152 | 3.599 | (8, -24, 33) | 0.04844 | (9, -43, 29) |
| | External capsule | L | 497 | 4.703 | (-31, -4, 18) | 0.05453 | (-31, -3, 18) |
| | | R | 393 | 4.336 | (29, -5, 18) | 0.05597 | (26, 17, 8) |
| | Fornix (cres) / Stria terminalis | L | 259 | 4.318 | (-25, -31, -4) | 0.06359 | (-25, -31, -4) |
| | | R | 177 | 4.662 | (26, -28, -5) | 0.07071 | (26, -28, -5) |
| | Genu of corpus callosum | | 4325 | 4.745 | (-14, 40, 2) | 0.05831 | (14, 36, -3) |
| | Posterior corona radiata | L | 1860 | 3.815 | (-27, -51, 27) | 0.04199 | (-27, -51, 27) |
| | | R | 2259 | 3.538 | (21, -28, 37) | 0.04046 | (20, -28, 37) |
| | Posterior limb of internal capsule | L | 279 | 4.051 | (-27, -10, 18) | 0.04858 | (-22, -24, 12) |
| | | R | 519 | 4.106 | (23, -22, 11) | 0.05517 | (23, -22, 11) |
| | Posterior thalamic radiation | L | 764 | 4.532 | (-31, -70, 9) | 0.0591 | (-31, -70, 9) |
| | | R | 1400 | 4.302 | (30, -72, 7) | 0.05847 | (30, -58, 12) |
| | Retrolenticular part of internal capsule | L | 1513 | 3.976 | (-33, -31, 0) | 0.04547 | (-36, -35, -3) |
| | | R | 1284 | 4.174 | (26, -23, -2) | 0.05305 | (26, -22, -3) |

| Category | ROI | Hemisphere | Voxel Count | Peak T | Peak T MNI coordinates | Peak Std. ß | Peak Std. ß MNI coordinates |
|---|---|---|---|---|---|---|---|
| | Sagittal stratum | L | 74 | 3.26 | (-37, -33, -4) | 0.0389 | (-37, -34, -4) |
| | | R | 1039 | 4.449 | (38, -39, -6) | 0.06798 | (38, -39, -6) |
| | Splenium of corpus callosum | | 4871 | 4.324 | (11, -43, 20) | 0.06404 | (11, -43, 12) |
| | Superior corona radiata | L | 6814 | 4.894 | (-29, -9, 21) | 0.05163 | (-30, -2, 20) |
| | | R | 6735 | 4.646 | (27, -5, 23) | 0.05791 | (16, 9, 39) |
| | Superior fronto-occipital fasciculus | L | 217 | 4.143 | (-22, 11, 23) | 0.04422 | (-20, 10, 24) |
| | | R | 165 | 4.199 | (22, 13, 23) | 0.04574 | (22, 14, 22) |
| | Superior longitudinal fasciculus | L | 4484 | 4.928 | (-31, -4, 20) | 0.06303 | (-32, -2, 18) |
| | | R | 4751 | 4.717 | (35, -6, 22) | 0.04913 | (35, -6, 22) |
| | Tapetum | L | 15 | 2.67 | (-26, -51, 20) | 0.0324 | (-24, -45, 21) |
| cerebellum/ brainstem | Cerebral peduncle | L | 44 | 3.199 | (-6, -22, -21) | 0.03881 | (-6, -22, -21) |
| | | R | 52 | 2.966 | (6, -21, -21) | 0.03431 | (9, -25, -17) |
| | Corticospinal tract | L | 1137 | 3.981 | (-7, -19, -29) | 0.05123 | (-7, -19, -29) |
| | | R | 1078 | 3.851 | (7, -24, -39) | 0.04962 | (7, -25, -40) |
| | Inferior cerebellar peduncle | L | 307 | 4.237 | (-8, -40, -40) | 0.05763 | (-8, -40, -40) |
| | | R | 293 | 3.531 | (10, -40, -37) | 0.04878 | (10, -40, -37) |
| | Medial lemniscus | L | 442 | 3.829 | (-5, -39, -40) | 0.05124 | (-7, -39, -40) |
| | | R | 466 | 3.887 | (8, -36, -28) | 0.05204 | (8, -37, -28) |
| | Middle cerebellar peduncle | - | 9882 | 4.885 | (29, -50, -43) | 0.07082 | (29, -50, -43) |
| | Pontine crossing tract (a part of MCP) | - | 1422 | 3.681 | (-6, -33, -36) | 0.04745 | (-7, -33, -37) |
| | Superior cerebellar peduncle | L | 118 | 3.291 | (-6, -31, -21) | 0.04009 | (-6, -31, -21) |

| Category | ROI | Hemisphere | Voxel Count | Peak T | Peak T MNI coordinates | Peak Std. ß | Peak Std. ß MNI coordinates |
|---|---|---|---|---|---|---|---|
| | | R | 116 | 3.332 | (5, -34, -21) | 0.04241 | (8, -37, -26) |
